# Walking along chromosomes with super-resolution imaging, contact maps, and integrative modeling

**DOI:** 10.1101/374058

**Authors:** Guy Nir, Irene Farabella, Cynthia Pérez Estrada, Carl G. Ebeling, Brian J. Beliveau, Hiroshi M. Sasaki, Soun H. Lee, Son C. Nguyen, Ruth B. McCole, Shyamtanu Chattoraj, Jelena Erceg, Jumana AlHaj Abed, Nuno M. C. Martins, Huy Q. Nguyen, Mohammed A. Hannan, Sheikh Russell, Neva C. Durand, Suhas S.P. Rao, Jocelyn Y. Kishi, Paula Soler-Vila, Michele Di Pierro, José N. Onuchic, Steven Callahan, John Schreiner, Jeff Stuckey, Peng Yin, Erez Lieberman Aiden, Marc A. Marti-Renom, C.-ting Wu

**Affiliations:** Department of Genetics, Harvard Medical School, 77 Avenue Louis Pasteur, Boston, MA 02115, USA.; CNAG-CRG, Centre for Genomic Regulation (CRG), Barcelona Institute of Science and Technology (BIST), Baldiri i Reixac 4, 08028, Barcelona, Spain.; Center for Genome Architecture, Department of Molecular and Human Genetics, Baylor College of Medicine, 1 Baylor Plaza, Houston, TX 77030, USA.; Center for Theoretical Biological Physics, Rice University, 6100 Main Street, Houston, TX 77005, USA.; Bruker Nano Inc., Suite 230, 630 Komas Drive, Salt Lake City, UT 84108, USA.; Wyss Institute for Biologically Inspired Engineering, Harvard University, 3 Blackfan Circle, Boston, MA 02115, USA.; Department of Systems Biology, Harvard Medical School, 200 Longwood Avenue, Boston, MA 02115, USA.; Broad Institute of the Massachusetts Institute of Technology and Harvard University, 415 Main Street, Cambridge, MA 02142, USA.; Department of Structural Biology, Stanford University School of Medicine, Stanford, CA 94305, USA.; Zero Epsilon, LLC, Suite 245, 699 East South Temple, Salt Lake City, UT 84102, USA.; Bruker Nano Inc., Suite 140, 3030 Laura Lane, Middleton, WI 53562, USA.; Departments of Computer Science and Computational and Applied Mathematics, Rice University, 6100 Main Street, Houston, TX 77005, USA.; Gene Regulation, Stem Cells and Cancer Program, Centre for Genomic Regulation (CRG), Dr. Aiguader 88, 08003, Barcelona, Spain.; Universitat Pompeu Fabra (UPF), Plaça de la Mercè 10, 08002, Barcelona, Spain.; ICREA, Pg. Lluís Companys 23, 08010, Barcelona, Spain.

## Abstract

Chromosome structure is thought to be crucial for proper functioning of the nucleus. Here, we present a method for visualizing chromosomal DNA at super-resolution and then integrating Hi-C data to produce three-dimensional models of chromosome organization. We begin by applying Oligopaint probes and the single-molecule localization microscopy methods of OligoSTORM and OligoDNA-PAINT to image 8 megabases of human chromosome 19, discovering that chromosomal regions contributing to compartments can form distinct structures. Intriguingly, our data also suggest that homologous maternal and paternal regions may be differentially organized. Finally, we integrate imaging data with Hi-C and restraint-based modeling using a method called *i*ntegrative *m*odeling of *g*enomic *r*egions (IMGR) to increase the genomic resolution of our traces to 10 kb.

**One Sentence Summary:** Super-resolution genome tracing, contact maps, and integrative modeling enable 10 kb resolution glimpses of chromosome folding.

## Main text

In terms of organization, chromosomal DNA is among the most challenging of biological molecules to study. It is massive, differs in sequence and structure from one region to the next, and often comes in pairs of homologs that defy easy distinction. Nevertheless, much progress has been made via genetic and epigenetic analyses, including DNA-DNA proximity ligation assays, such as Hi-C and other Chromosome Conformation Capture technologies (1–8), as well as three-dimensional (3D) modeling (reviewed by (9, 10)). Recently, researchers have walked along chromosomes using widefield microscopy and fluorescent *in situ* hybridization (FISH) probes in rounds of hybridization (11), wherein each round targeted the central portion of a Hi-C defined self-interacting contact domain, also known as a topologically associating domain, or TAD (3–7). This strategy revealed the overall path of chromosomes, which was then used as a guideline for Hi-C contact-based modeling (bioRxiv, https://doi.org/10.1101/318493 May 2018). Other studies, using super-resolution microscopy, have explored a multitude of genomic features (12–18), some lending support for a physical basis for contact domains (15, 18). What remains relatively unexplored, is the physical nature of compartmental intervals, that is, the contiguous chromosomal regions whose loci share long-range contact patterns and collectively, generate the plaid “compartment” pattern often seen extending above and below the Hi-C diagonal (3).

Here, we present a method for visualizing chromosomal regions at super-resolution and then integrating the images with Hi-C interaction matrices via a data-driven computational protocol to produce 3D models of chromosome organization at the level of the single cell. We begin by describing our strategy for walking along 8.16 megabases (Mbs) of human chromosome 19 in primary skin fibroblasts of donor PGP1 (XY, PGP1f; Methods), using oligonucleotide-based Oligopaint FISH probes (19, 20) and imaging in sequential steps using the single-molecule localization microscopy methods of OligoSTORM and OligoDNA-PAINT (12, 20). The resulting images successfully resolved the structure of individual compartmental intervals, showing further that it can vary from cell to cell. We then describe a strategy for integrating images of the genome with Hi-C contact frequency data to produce 3D models at 10 kilobase (kb) resolution. As the PGP1 genome has been haplotyped and its pedigree is known, (Fig. 1A, B; (21); Methods) we also describe a strategy whereby the maternal and paternal homologs can be distinguished.

**Fig. 1.**
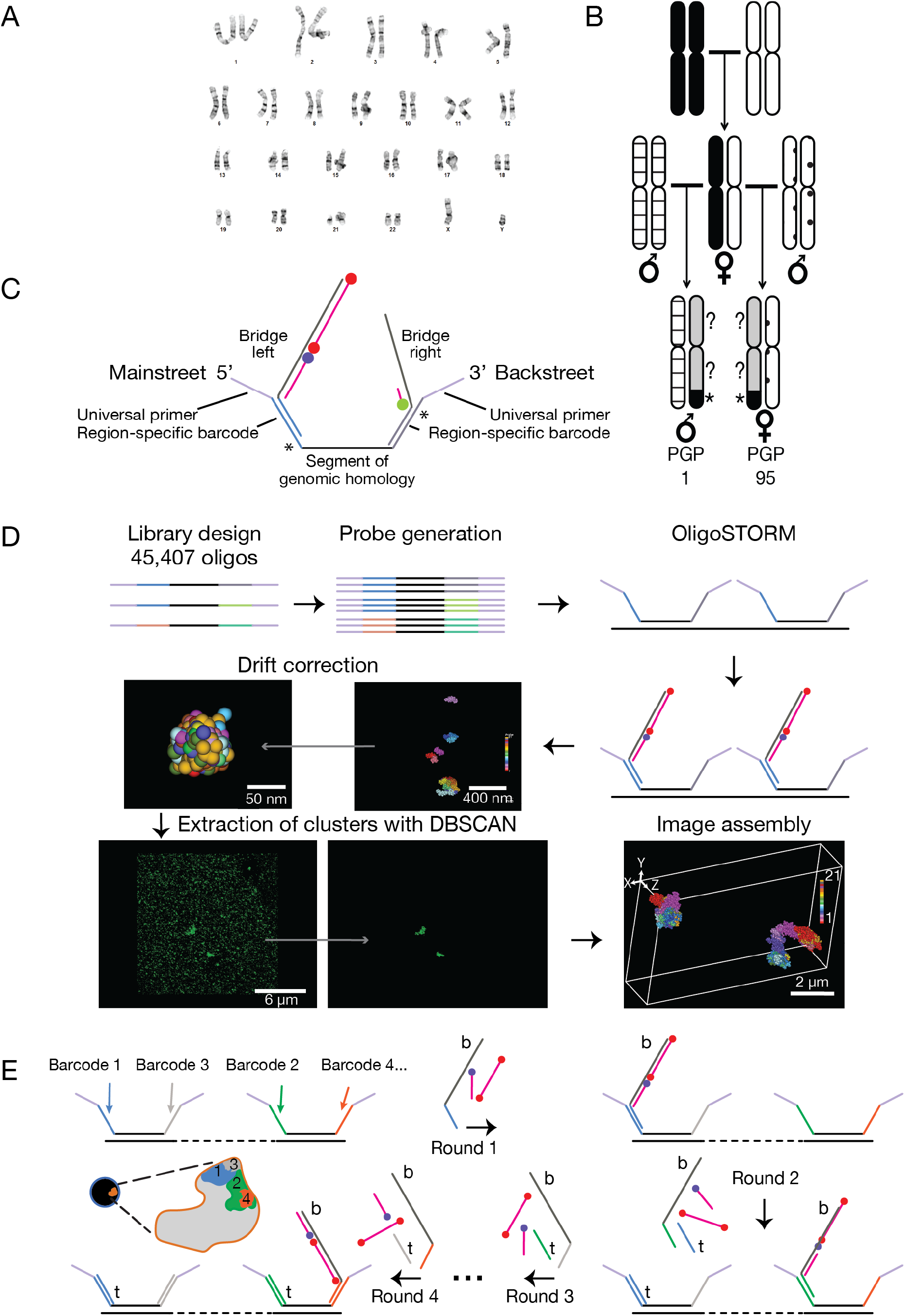
Tools. (**A**) PGP1f karyotype (Methods). (**B**) Assignment of parental origin to homologs of chromosome 19 through pedigree analysis. Black and white, homologs of chromosome 19 in maternal parent of PGP1 and PGP95; patterned, homologs of paternal parents; asterisk and black, SNV block shared between PGP1 and PGP95 identifying maternal homolog; ‘?’ and grey, maternally derived regions not shared by PGP1 and PGP95. (**C**) Oligopaint oligos (129 −135 nts) consist of a segment of genomic homology (black; 35 - 41 nts) flanked by Mainstreet and Backstreet (47 nts), each consisting of universal primers (violet; 20 nts), region-specific barcodes (blue and grey; 20 nt), and a 7-nt sequence (asterisk) which, in combination with barcodes, generates a binding site for toehold oligos. A bridge oligo (left) bound to a Mainstreet barcode allows labeled secondary oligos (blue, Alexa Fluor 405; red, Alexa Fluor 647) to co-localize and enable OligoSTORM. A bridge (right) bound to a Backstreet barcode allows a labeled imager strand (green, Cy3B) to bind transiently, enabling OligoDNA-PAINT. (**D**) Workflow from library design through probe generation, OligoSTORM, drift correction using 90 nm fiducial markers, extraction of clusters with DBSCAN, and image assembly, as described in the text. (**E**) Walking along chromosomes using sequential hybridizations. All Oligopaint oligos are hybridized to the genome at once, and then rounds of hybridization and imaging are conducted. Each round brings in bridges (‘b’), labeled secondary oligos, and, except for round 1, toehold oligos (‘t’) in order to label a new region of the walk while removing labeled oligos from the previously imaged region.

We considered two approaches for walking along the chromosome – one in which the sizes of the steps are in accordance with genomic features and another in which step sizes are uniform – initiating our studies with the first. Thus, we designed Oligopaint probes corresponding to the Hi-C defined features of compartments as well as smaller features, including contact domains, loops, and genes. Because a Hi-C map of PGP1f cells was not available when we initiated our study, we designed our walk based on the Hi-C data of IMR-90 lung fibroblasts (Fig. S1; (8)) Methods). Focusing on an 8.16 Mb region extending from chr19:7,400,000 to chr19:15,560,000, we then generated Oligopaint probes to image the region in 9 segments ranging in size from 360 kb to 1.8 Mb, including probes that would permit visualization of features as small as 10 kb (Table S1; also see Bintu, *et al*.).

Similar to fluorophore- (dye-) conjugated oligonucleotides (oligos) pioneered over two decades ago (22) as well as other iterations (Methods), Oligopaint probes consist of computationally designed oligos (Fig. 1C; (19)). In addition, they are single-stranded and carry a short region (~32-42 nucleotides, nts) of genomic homology as well as nongenomic sequences, called Mainstreet and Backstreet, at their 5’ and 3’ ends, respectively. Streets permit Oligopaints to be amplified, multiplexed, barcoded, and indirectly visualized via labeled secondary oligos (11, 12, 14, 19, 20, 23–26). Here, we use Streets to enable OligoSTORM and OligoDNA-PAINT (12, 14, 20). OligoSTORM melds Oligopaints with STORM (Stochastic Optical Reconstruction Microscopy; (27)) or dSTORM (direct STORM; (28)) and generates single events of fluorescence (localizations) via single dyes or dye-pairs. As for Oligo-DNA-PAINT, it combines Oligopaints with DNA-PAINT (DNA-points accumulation for imaging in nanoscale topography; (29, 30)) and generates localizations via transient hybridizations of short (~9-10 nts) dye-labeled oligos, called imager strands, to complementary sequences, called DNA docking sites, located on the Streets; here the transient binding of imager strands is interpreted as localizations.

Consisting of 45,407 species of oligos, our Oligopaint library incorporated three key features (Fig. 1C). First, all the oligos avoid sequences that differ between the parental genomes of PGP1f and, thus, they bind both homologs of chromosome 19. Second, both streets of each oligo carry 20-nt long barcodes corresponding to sub-regions of our 8.16 Mb walk; these barcodes confer multiple functionalities on the oligo. For example, when enabling OligoSTORM, these barcodes co-localize “bridge” oligos that recruit the labeled secondary oligos to be visualized by OligoSTORM. When enabling OligoDNA-PAINT, the bridges include a docking site for the imager strand. The barcodes also permit any targeted genomic region to be imaged twice, once via Mainstreet and then, again, via Backstreet. Finally, each oligo carries a 7-nt sequence (Fig. 1C, asterisk) that facilitates sequential rounds of imaging; in conjunction with a flanking barcode, these 7 nts generate a region which, when bound by a 27-nt “toehold” oligo (31–33), dislodge any previously bound bridge.

Our workflow (Fig. 1D; Methods) begins with T7 RNA polymerase-mediated amplification of the entirety of an Oligopaint library that is to be used for a walk. The totality of the resulting single-stranded oligos (11) is then hybridized to fixed cells in a single round of denaturation and hybridization. When implementing OligoSTORM, the sample is then placed on a 3D single-molecule localization microscope (Bruker Vutara 352) and scanned to identify nuclei to be imaged. Sequential rounds of imaging are then conducted with an automated microfluidic system (Fig. 1E), each round targeting one step of the walk by hybridizing a specific bridge oligo (Fig. 1E, labeled with ‘b’) along with its dye-conjugated oligos, after which images are acquired one nucleus at a time until all nuclei have been imaged; the next round is initiated through the use of toehold oligos (Fig. 1E, labeled with ‘t’), which dislodge bridges bound in the previous round. To traverse the depth of PGP1f nuclei, imaging is conducted in 100 nm increments along the axial (Z) dimension for up to 4 *μ*m, well within the volumetric capability of the Vutara. This protocol has given an average localization precision of 11.14 +/− 1.47 nm and 11.02 +/− 1.22 nm along the lateral axes and 46.96 +/− 6.02 nm along the axial dimension, corresponding to average supported optical resolutions of 26 ± 3 nm and 112 ± 11 nm, respectively. All localizations are then subjected to drift correction with respect to fiducial beads, after which density-based spatial clustering of applications with noise (DBSCAN; (34)) extracts clusters of localizations most likely to represent genomic regions (Fig. 1E; Methods).

Using our workflow, we have walked the entirety of the 8.16 Mb region in over 60 chromosomes, separately imaging all 9 chromosomal segments via Mainstreet (CS1-9; Fig. 2A, B; Movie S1; Table S1). These studies also demonstrated the effectiveness of using Backstreet in conjunction with Mainstreet to image smaller features lying within larger regions, all in one imaging run (Table S1); for example, in a single 21-round imaging run that imaged CS1-9 using Mainstreet (Fig. 2A, rounds 1-9), we used Backstreet and to image four subregions comprising CS7 (Fig. 2A, rounds 10-13; Fig. S2), a loop, including its anchors and flanking regions in CS6 (Fig. 2C, rounds 14-18), and the Dnmt1 gene in CS3 (Fig. 2D, round 19). As a further demonstration of the capacity of barcodes, we have taken advantage of the propensity of chromosomes to lie within their own territories (35) and traced several chromosomes in parallel. For example, we have traced the 8.16 Mb region of chromosomes 19 while also walking along chromosomes 3 and 5 in uniform step sizes of 500 and 250 kb, respectively (Fig. 2E; Fig. S3 for all 23 steps; Table S2; Methods). We have also used our library to image CS7, 8, and 9 using OligoDNA-PAINT (Fig. 2F; Table S3), achieving an exciting supported resolution of 18 nm (Methods; (36)). Finally, to assess the reproducibility of our images, we have conducted a number of assessments using OligoSTORM (Methods). In one, we imaged CS1 using Mainstreet and then, imaged it again, 90 hours later using Backstreet, achieving an average spatial overlap of 80 ± 9% (n = 20; Methods) with minimal change in the average number of localizations (n=4,8371 for round 1 and 4,912 for round 21; Fig. 2G, S4). Comparable results were obtained (75% - 85%) when we imaged CS7 via Mainstreet and then reimaged it using barcodes on Backstreet to sequentially visualize four subregions comprising CS7 (Fig. 2H).

**Fig. 2.**
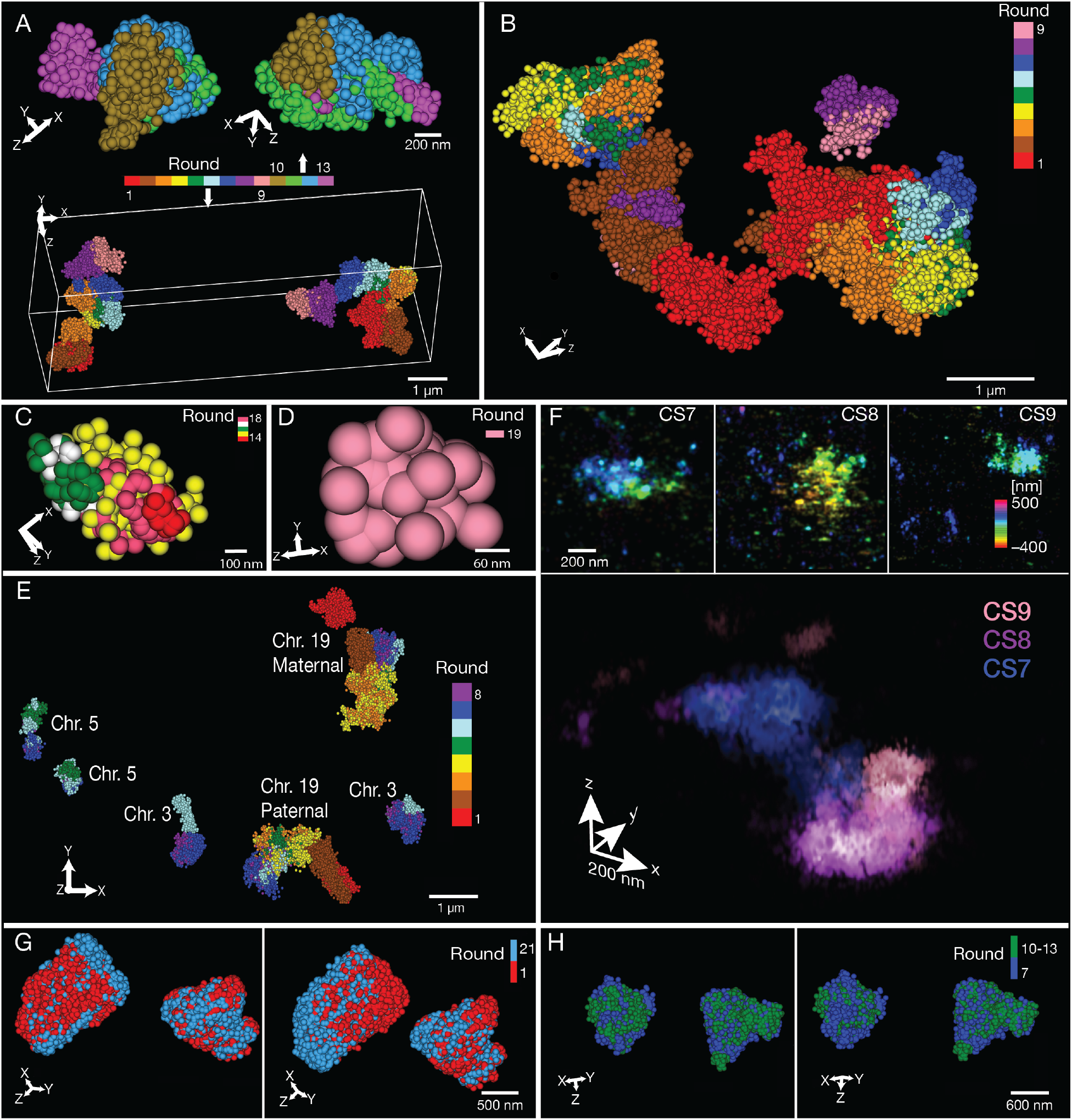
Tracing PGP1f chromosomes. (**A**) OligoSTORM image of a diploid nucleus showing CS1-9 (Bottom, rounds 1 through 9; 1.28, 1.24, 1.80, 1.04, 0.56, 0.52, 0.84, 0.52, and 0.36 Mb, respectively) and four subregions of CS7 (top, rounds 10 through 13; 140, 260, 350, and 90 kb, respectively) of both homologs. In this and all panels excepting F, radius of spheres represents the localization precision in the axial dimension. (**B**) Same as in A with respect to CS1-9, but a different nucleus. (**C**) Loop. Red and green, loop anchors (10 kb each); yellow, loop body (270 kb); pink and white, flanking regions (20 and 80 kb, respectively). (**D**) *DNMT1* (59.5 kb). (**E**) Walking along chromosome 19 in variable step sizes (see A) while also walking along chromosomes 3 and 5 in uniform step sizes (500 and 250 kb, respectively). (**F**) OligoDNA-PAINT images of CS7-9 of one homolog showing each chromosomal segment by itself (top; color scale represents depth in z) or merged (bottom; color scale represents different chromosomal segments). In contrast to other panels, localizations are blurred according to precision. (**G**) Superimposition of two images, taken 90 hours apart, each of both homologs of CS1 in (red, first image; blue, later image). Average overlap is 80 ± 9% and 68 ± 12% (n = 20) when images are aligned based on their centers of mass (left) or not (right), respectively. (**H**) Superimposition of two images from one nucleus of both homologs of CS7, one image encompassing all of CS7 (blue) and the other being a composite of the four subregions (green; 140, 260, 350, and 90 kb) comprising CS7. With alignment based on centers of mass, 85% of the single image overlapped the composite image, and 76% of the latter overlapped the former (left). Without alignment, the analogous values were 84% and 75% (right). Note, the coordinates of the four subregions were based on the IMR-90 Hi-C map to correspond to contact domains. Sequencing depth of the PGP1f Hi-C map was not able to confirm these as contact domains in PGP1f, and thus they are referred to simply as subregions.

Intriguingly, we noticed early on that a number of consecutive steps of our walk appeared minimally entangled, suggesting that they correspond to distinct physical entities. To explore this, we further analyzed 38 OligoSTORM walks of CS1-9; these 38 represented the 19 nuclei with the highest image quality and for which we observed exactly two foci of signal per chromosomal segment and no evidence of aneuploidy (Methods). These nuclei were analyzed by convolving them with a Gaussian kernel (37), which enabled us to represent each chromosomal segment of each chromosome as a distribution of localization densities in 3D space (Methods). To assess entanglement of chromosomal segments, we used our density maps and calculated centers of mass, distances between centers of mass (Distance score, DS) and overlap (Entanglement score, ES; Fig. S5; Methods). The results revealed that chromosomal segments are not randomly placed with respect to each other. Although there was considerable variation in DS (Fig. S5B), the observed DS distributions and their mean distance scores (MDS) showed that the five central segments, CS3-7, were closer than expected in all pairwise combinations (Fig. 3A) except one (CS4 and CS7), which nevertheless trended in the same direction (Fig. S5B). Overall, these central segments were also distanced from the segments lying at the beginning (CS1 and CS2) and end (CS8 and CS9) of the walk. Importantly, even with only 38 walks, these differences from expectation were statistically significant for most pairwise comparisons (Fig. S5B). Moreover, segments at the edges of the central region were further than expected from segments just beyond the central region; CS3 was further from CS2, and CS7 was further from CS8 than expected. Interestingly, the beginning (CS1 and CS2) and end (CS8 and CS9) of the walk trended towards being closer than expected. Altogether, these observations suggested the center of the walk to be spatially separated from the ends, with the two ends being closer in space than expected.

**Fig. 3.**
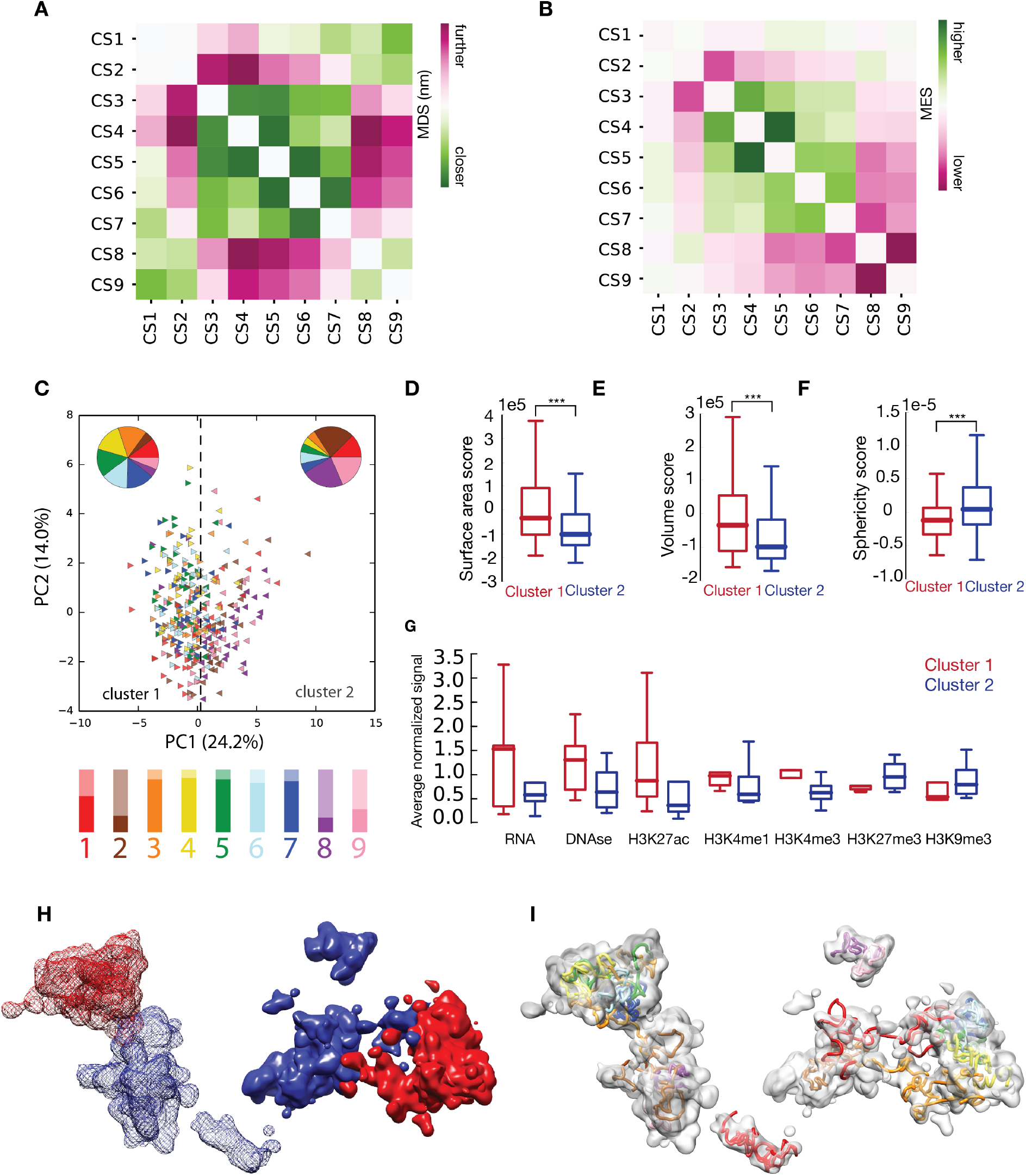
Image analysis and integrative modeling. (**A, B**) Matrices for (**A**) mean spatial distance scores (MDS) and (**B**) mean entanglement scores (MES) for CS1-9. (**C**) Unsupervised PCA of 3D features for the 342 visualized chromosomal segments suggests two major clusters. Chromosomal segments are colored as in Figures 2A and B, with relative abundances displayed as pie charts. Lower panel, proportion of images of each chromosomal segment in cluster 1 (opaque color) or cluster 2 (transparent color). (**D-F**) Variation and distribution of the two clusters in terms of (**D**) surface area, (**E**) volume, and (**F**) sphericity scores are shown as box plots, where boundaries represent 1st and 3rd quartiles, middle line represents median, and whiskers extend to 1.5 times the interquartile range (Mann-Whitney rank test, ***: p < 10^−3^). (**G**) Mean normalized signal of seven different ChIP-seq profiles in PGP1 nuclei for the two identified clusters. Red, cluster 1; blue, cluster 2. (**H**) Density map representation of each chromosomal segment in both copies of the entire 8.16 Mb (CS1-9) region in a single PGP1 nucleus as imaged by OligoSTORM, color-coded by cluster type. Red, cluster 1; blue, cluster 2. One homolog is displayed with a solid surface, while the other homolog is displayed with a mesh surface. Note spatial segregation of clusters. (**I**) Density map representation of CS1-9 as in I (grey) except displayed, here, with the fitted 3D model obtained via IMGR (colored coded as in Fig. 2A, B). Images were created with Chimera (50).

Our analyses of entanglement also revealed a nonrandom pattern (Fig. 3B, S5C, D; Methods (37)). Although, again, there was considerable variation (Fig. S5D), the distributions of ES and their mean entanglement scores (MES) revealed that the central segments, CS3-7, were more entangled than expected or trended as such, while the segments lying at the boundaries of this region (CS3 and CS7) entangled less than expected with the segments lying upstream and downstream, respectively (Figs. 3B, S5D). The contiguous pairs of segments lying at the ends of the walk were also minimally entangled; CS1 and CS2 showed insignificant entanglement, while CS8 and CS9 resulted in the lowest of all the MES scores. Thus, it is especially noteworthy that CS1 and CS2, lying at one end of the walk, nevertheless showed significant entanglement with CS9 and CS8, respectively, at the other end (Figs. 3B, S5D). These observations aligned with our analyses of distance, suggesting a 3D clustering of the 9 chromosomal segments into two groups, one consisting of the central segments (CS3-7) and the other consisting of the ends of the walk (CS1, 2, 8, and 9). This was confirmed by unsupervised clustering of the MDS and MES matrices (Fig. S5E, F).

To further characterize the clustering, we calculated the surface area, volume, and sphericity for each chromosomal segment, corrected for genomic size (Fig. S6; Methods) and then, combining these features with the 8 measures of distance (DS) and 8 measures of entanglement (ES), conducted unsupervised clustering via Principal Component Analyses (PCA) for all 342 imaged chromosomal segments (38 chromosomes x 9 chromosomal segments) (Methods; Fig. S7). Two major clusters emerged, with the first two principal components accounting for 24.2% and 14.0% of the variability (Fig. 3C, S7A); the first cluster comprised 73.8% of all the images of CS3-7, while the second cluster comprised 75.0% of all the images of CS1, 2, 8 and 9. Considered together, the segments of cluster 1 had larger area and volume and were less spherical than the aggregate of the segments in cluster 2 (Fig. 3D-F). Interestingly, the segments that were primarily in cluster 1 harbored epigenetic markings of active chromatin, while those primarily in cluster 2 harbored markings of inactive chromatin (Fig. 3G; Methods). These findings align with the differing degrees of chromatin compactness that have been observed for different epigenetic states in *Drosophila* (14).

Our analyses also highlighted variability in the classifications for all chromosomal segments (Table S4). For example, while CS3-7 fell primarily in cluster 1 and CS2 and CS8 fell primarily in cluster 2, CS1 and CS9 resulted in much more mixed classifications, with ratios of cluster 1:cluster 2 being 42:58 and 37:63, respectively (Fig. 3C, bar graph). CS1 is particularly noteworthy in this regard; the distributions of PC1 values revealed that all segments were skewed towards one cluster type with the exception of CS1 (Fig. S7B-J), which was also the most mixed in terms of intra-nuclear and inter-nuclear variation (Fig. S7K). In brief, our data suggest that, while chromosomal segments can be broadly classified by their structural properties into categories reminiscent of active and/or inactive chromatin, they are also variable and dynamic in their character.

In sum, OligoSTORM derived density maps suggested a number of organizational traits for the imaged region (Fig. 3H). First, CS3-7, can form an internally entangled entity. Second, this region can be distanced from, as well as less entangled with, its flanking segments, suggesting the potential of distance, *per se*, to be an organizational principle. Third, the entire region can fold back on itself such that its ends come into proximity and entangle to a modest degree (see example in Fig. 2B). Fourth, CS1, CS2, CS8, and CS9 may each correspond to an individual entity, suggesting that the imaged region may be composed of at least 5 structurally distinct sections. Fifth, the two clusters of segments revealed by PCA may correspond to spatially separated compartments, active (A-type) and relatively inactive (B-type), reminiscent of those observed by Hi-C (3). Finally, while some chromosomal segments were predominantly active or inactive, others could be classified as in either state or mixed.

How well might our images correlate with population-based data from DNA-DNA proximity ligation based technologies? To address this, we generated an *in situ* Hi-C map for PGP1f cells (Fig. 4; Table S5; Methods) and assigned loci in the target region to A and B compartments based on the resulting map. The annotation assigned CS1, CS3-7, and CS9 to the A compartment, whereas CS8 and most of CS2 were assigned to the B compartment (Fig. 4, S8). Notably, the fact that CS3-7 fell in a single compartment meant that all loci in this extended interval exhibited enhanced contact frequencies with one another. These findings recapitulate several features of the imaging data: [i] five physical entities (CS1, CS2, CS3-7, CS8, and CS9) with distinctive chromatin states; [ii] a single, extended structure spanning CS3-7; [iii] elevated contact frequency between the endpoints of the target region; and [iv] strong correspondence between the chromosomal segments assigned to the A compartment (CS1, CS3-7, and CS9) and those that were frequently assigned to cluster 1 (CS1and CS3-7). This strong correspondence between the outcomes of two very different technologies, OligoSTORM and Hi-C, lends confidence to our image-based measurements of distance, entanglement, size, volume, sphericity, and structural variability as well as to the overall potential of OligoSTORM to elucidate the organization of large swaths of, if not entire, chromosomes. Moreover, our images specifically argue that compartmental intervals have the capacity to form distinct structures.

**Fig. 4.**
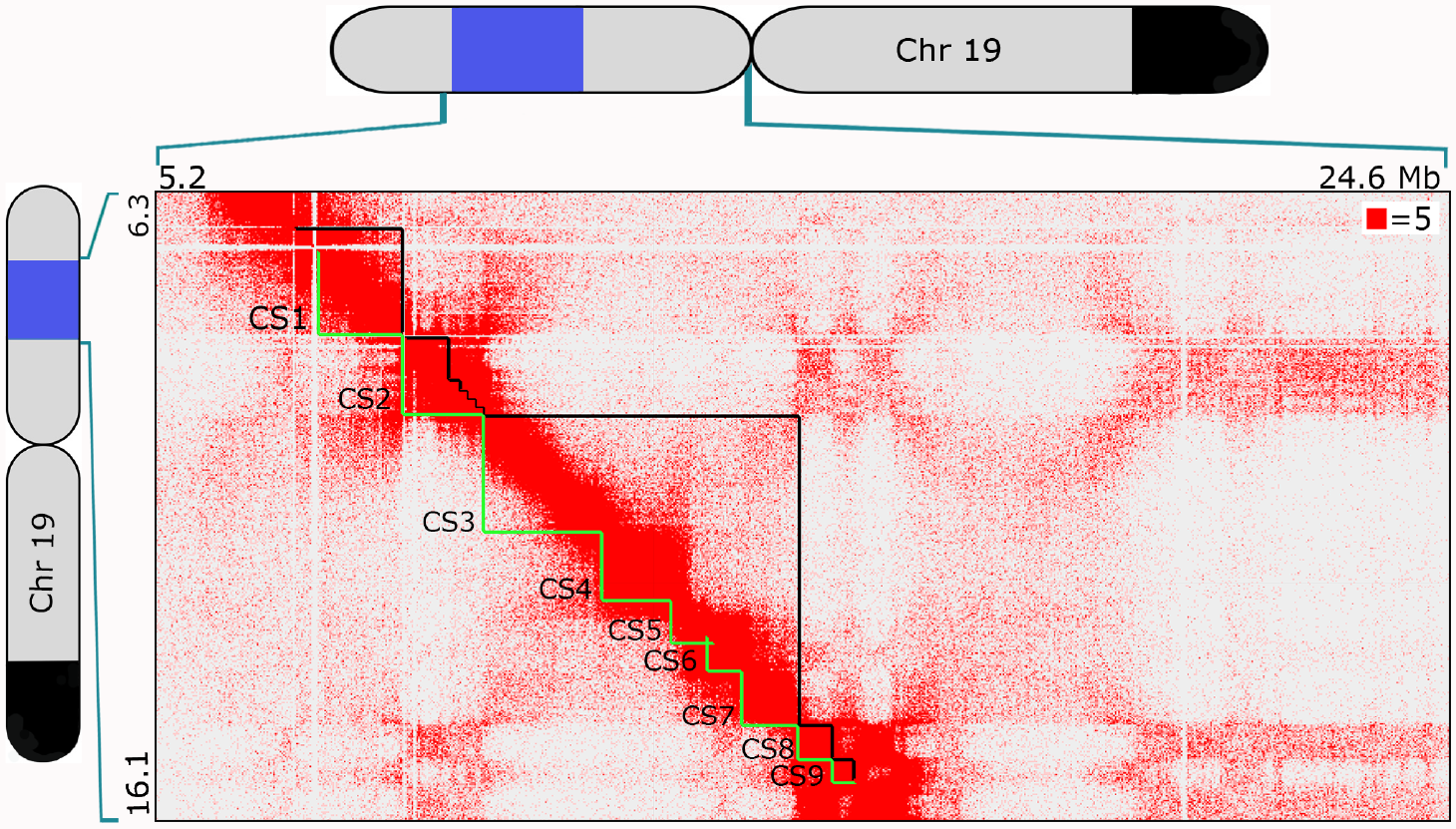
*In situ* Hi-C map of CS1-9 in PGP1f. Hi-C map of region including CS1-9, displayed at 25 kb resolution and normalized according to Knight & Ruiz (51). Contact count indicated by the color of each pixel, ranging from 0 (white) to 6 (red); OligoSTORM steps (CS1-9) delineated in green below the diagonal, and compartmental intervals delineated in black above the diagonal. Cartoons of chromosome 19 show extent of the 8.16 Mb imaged region (blue) and block of SNVs shared between PGP1 and PGP95 (black; as in Fig. 1B).

Next, we developed an integrative protocol for combining different modalities of data to model the genome at unprecedented genomic resolution. This integrative modeling of genomic regions (IMGR) was used here to integrate OligoSTORM and Hi-C data, producing, for the first time, a 3D model of two homologous regions in a single diploid nucleus (Figs. 3I, S9A; Movie S2; see Methods and Supplementary notes for rationale of IMGR, including rigid and flexible fittings and protocol to select models most likely to represent the image in question.). Population-based 3D modeling of Hi-C data has been used before to recapitulate diploid genomes (38), and 3D modeling of single-cell Hi-C data has been implemented for haploid reconstruction (39, 40). However, modeling homologous chromosomes has proven challenging due to structural variability between the two chromosome copies (39). In this study, we addressed diploidy through integrative modeling. We used, as input, an ensemble of Hi-C derived 3D models and the corresponding single-nuclei OligoSTORM density maps for each of the 9 chromosomal segments we had imaged; the models had been built as previously described (41, 42), using 10 kb-resolution PGP1f Hi-C maps. Then, via a protocol inspired by the fitting of proteins to cryoEM density maps (43), we implemented a two-step integrative process to fit each of the ensemble 3D models into the density map of the corresponding chromosomal segment (Fig. S9A; Methods). Thus, this modeling protocol achieved a resolution of 10 kb for a genomic region that had been imaged in step sizes ranging from 0.36 to 1.8 Mbs in size.

Models of the genomic resolution we have produced can yield insights into the 3D organization of specific loci in the modeled region. For example, chromosome 19 is extraordinarily rich in zinc-finger genes, which are clustered in 6 locations (44), and Figure S9B models the arrangement of the two clusters located in the region we imaged; one lies at one end of CS2 and another extends across CS4. Similarly, all genes in the imaged region that are classified by OMIM as disease-causing (45) can now be positioned in 3D (Fig. S9C). The models can also be used to map ChIP-seq data in 3D space (46). For instance, it appears that one end of each copy of the region is decorated by a large patch of H3K4me1, H3K27ac, and H3K4me3 (Fig. S9D-F). These patches suggest enhancer-promoter clusters and, consistent with this, RNA-seq and DNAse accessibility data indicate that the patches co-localized with foci of expressed genes and greater accessibility (Fig. S9G, H). In contrast, ChIP-seq data for facultative (H3K27me3) and constitutive (H3K9me3) heterochromatin suggest multi-foci patches of heterochromatin throughout (Fig. S9I, J).

Finally, we have explored the potential of our approach to address questions regarding the relationship between homologous chromosomal regions. Indeed, although not identified by haplotype, allelic differences in accessibility and volume have been noted at the level of individual genes (47). Will this difference be maintained across large chromosomal regions and, if so, will they reflect parental origin? Here, we assessed the difference between homologous regions by first assigning an ellipticity score to each 8.16 Mb region of both homologs for all 19 nuclei and then, for each nucleus, taking the ratio of the larger to the smaller score to obtain an ellipticity ratio (Methods). Intriguingly, the median ellipticity ratio was 2.5 (±0.54), which was significantly greater than for randomly selected pairs of homologous regions (1.3 ± 0.48; Fig. 5A, S10A, Table S6), suggesting that homologous regions of a single nucleus may differ nonrandomly in ellipticity and that the difference is, on average, greater between homologous regions within a nucleus than between nuclei.

**Fig. 5.**
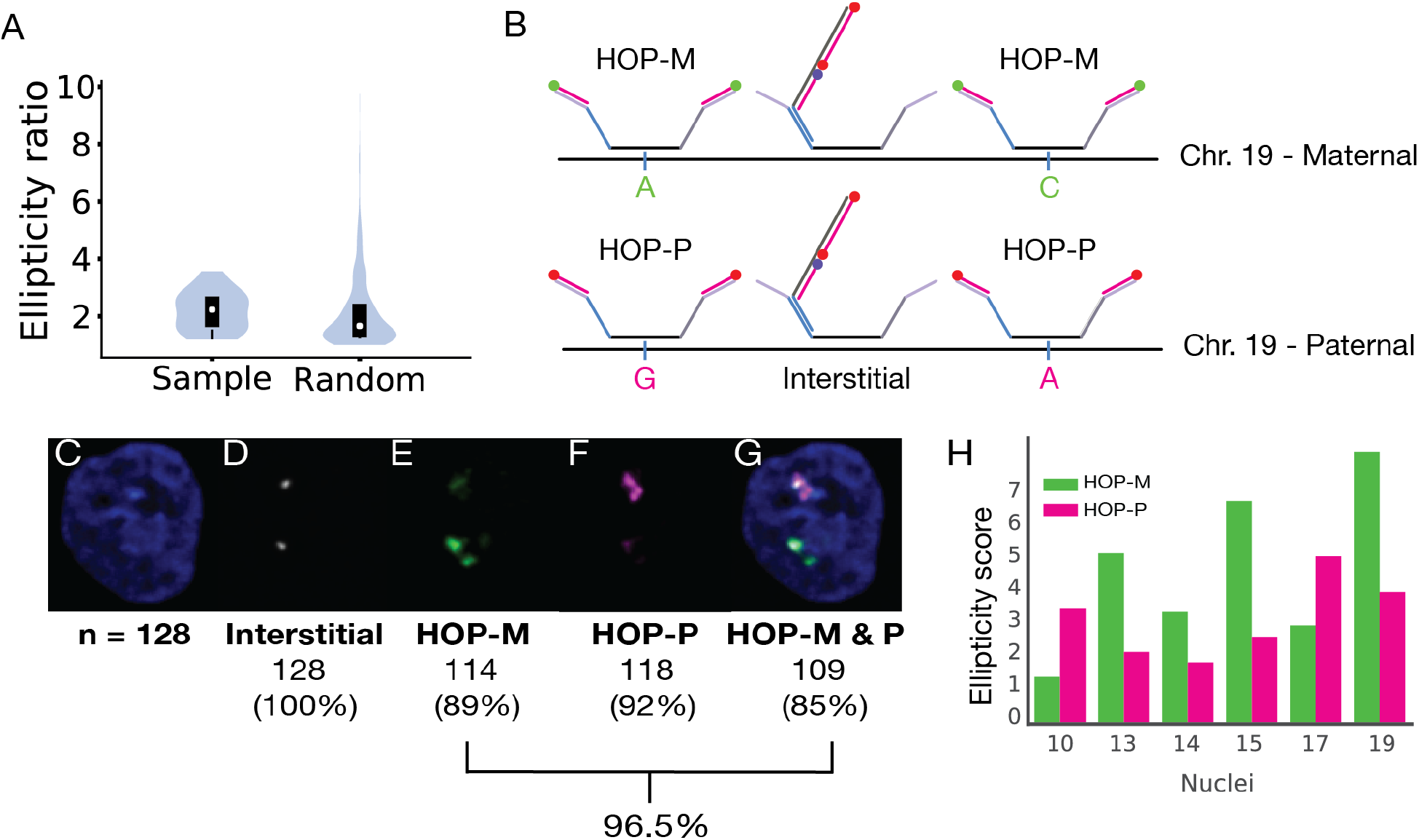
Homolog-specific Oligopaints for PGP1. (**A**) Violin plots of the ratios of 19 pairs of ellipticity scores, each pair representing the two homologs of one of the 19 imaged nuclei (Sample), and of 1,000 pairs of ellipticity scores, each pair representing two chromosomes chosen at random from the 38 representing the 19 imaged nuclei (Random). Boundaries of the black box-plot represent 1st and 3rd quartiles, white dot represents the median, and whiskers extend to 1.5 times the interquartile range (Mann-Whitney rank test, *: p < 10^−2^). (**B**) The HOP-M and HOP-P probes for chromosome 19 each encompass the entire chromosome and contain 11,259 oligos that cover ≥ 1 SNV per oligo. Thus, they are in contrast to Oligopaint oligos used to image CS1-9, these latter probes being “interstitial” in nature, as they avoid SNVs. HOP-M and HOP-P probes are visualized with secondary oligos labeled with different dyes such that the two probes can be distinguished. (**C-G**) Images of a nucleus visualized with DAPI (**C**, blue), an interstitial probe targeting just CS3 (**D**, grey), HOP-M (**E**, green, Atto488N), HOP-P (**F**, magenta, Atto565N), and all probes (**G**), the latter demonstrating co-localization of all signals (n = 128); percentages show efficiency of each probe configuration, with a combined efficiency of 96.5%. (**H**) Ellipticity for the maternal (green) and paternal (magenta) homologs of the 6 nuclei for which HOPs had been applied.

To ascertain whether greater ellipticity might correlate with parental origin, we turned to the most recent of our 38 walks as, for these, we had been successful in assigning parental origin. Here, we had implemented homolog-specific Oligopaints (HOPs; (12)), which target single nucleotide variants (SNVs) that differ between the haplotypes of a genome and, thus, permit distinction of homologous genomic regions. While proven successful in *Drosophila* and mice with abundant SNVs (~2-5 SNVs per kb; (12)), their applicability to humans, which have a lower frequency of SNVs (~0.77 SNVs per kb for PGP1), was unknown. Thus, we generated two libraries, HOP-M and HOP-P, the former targeting 11,259 maternal-specific SNVs extending across the entirety of chromosome 19, and the latter targeting the analogous 11,259 paternal-specific SNVs (Fig. 5B; Methods). Excitingly, HOP-M and HOP-P proved successful in differentially labeling the homologs (Fig. 5C-G, S10B, C). Thus, having applied HOPs in our most recent imaging runs, we were able analyze 12 walks (6 nuclei) for which parental origin was determined and conclude that ellipticity was not absolutely correlated with parental origin (Fig. 5H). Interestingly, MDS and MES matrices suggest differences between maternal and paternal homologs, with the potential of CS3-7 being more spatially compartmentalized in maternal as versus paternal homologs (Fig. S10D-I); of note, maternal and paternal homologs produced matrices reminiscent of those of the entire set of 38 chromosomes, arguing that the homologs we distinguished by HOPs approximate those of the larger population (Figs. 3A, B, S10F, I). While these studies must be greatly expanded to substantiate any homolog-specific trend, our experience indicates that OligoSTORM should be useful for exploring the biology of homology.

In conclusion, we have described strategies for *in situ* super-resolution genome visualization through sequential rounds of imaging, followed by integration of the resulting images with Hi-C data to produce 3D models at 10 kb resolution in single nuclei. In conjunction with other efforts (e.g., refs (11–18)), these strategies should advance our understanding of genome organization and, when enhanced with HOPs, the impact of parental origin. Indeed, here we have observed patterns of chromosome organization that were otherwise unknown. In line with other studies, including a live-cell analysis correlating chromatin accessibility more with fluctuation than compaction (bioRxiv, https://www.biorxiv.org/content/early/2018/07/09/365700 July 2018), we have also observed significant structural variability and speculate, here, as to the potential of that variability. Such variability may reflect genomic function or may, itself, be a signature that distinguishes one genomic region, or chromatin state, from another. For example, while regulatory factors may recognize their binding regions by the conformation of their targets, it may be that some also, or solely, respond to the variations of that conformation, the number of conformations, and/or the speed with which their targets transition through conformations. If propagated along chromosomes or of sufficient magnitude to generate a turbulence that can be propagated through the nuclear milieu, dynamic variability of chromosomal regions may even function to convey information across nuclear distances, perhaps even perturb or promote phase separations (48, 49). Finally, we note that, as our strategies are applicable to entire chromosomes, it should be possible to implement them in studies of chromosomes as single, fully integrated units of structure and function, consistent with the unit of inheritance being as much the chromosome as it is the gene.

## Acknowledgments

We thank Geoffrey Fudenberg for calling contact domains and compartmental intervals in IMR-90 cells; Kun Zhang, Daniel Jacobsen, Bing Ren, and Anthony Schmitt for PGP1 haplotyping; Bogdan Bintu for help regarding microfluidics; Bogdan Bintu, Alistair Boettiger, Florian Schueder, and Xiaowei Zhuang for helpful input; and Jennifer Waters and the Nikon Imaging Center of Harvard Medical School. The authors also thank members of the Yin, Aiden, Marti-Renom, and Wu laboratories for countless valuable discussions. Finally, C.-t.W. thanks H.N., in particular, for conversations regarding the implications of structural variation and turbulence and, for remarkable cells and years of discussion, she also thanks PGP1 and PGP95.

## Author contributions

G.N. conceived the project, led the design, performed experiments and analyses, and wrote the paper. I.F. conceived and performed analyses and integrative modeling and wrote the paper. C.P.E conceived and performed Hi-C experiments and analyses and wrote the paper. C.G.E conceived innovations to STORM imaging and data storage and analysis and wrote the paper. B.J.B, H.M.S., S.H.L., S.C.N, and R.B.M conceived several aspects of the project, performed experiments and analyses, and edited the paper. S.C., M.H., S.R., and N.C.D performed experiments. S.S.P.R, J.Y.K., P.S-V, and M.D.P. analyzed data. J.E., J.A.A, N.M.C.M, and H.Q.N contributed to discussions. M.D.P and J.N.O. contributed to Hi-C discussions. S.Ca and J.Sc conceived the Bruker Vutara image analysis software. J.St, P.Y., E.L.A, M.A.M-R, and C.-t.W participated in all aspects of the study and wrote the paper.

## Data and materials availability

All data from the study are either in the manuscript, available upon request, or as follows. Streets and MATLAB code used for the generation and appending of streets can be downloaded at https://github.com/gnir/OligoLego. PGP1f cells are available from the Personal Genome Project (https://www.personalgenomes.org/us) upon acceptance of a standard MTA. Hi-C data will be placed in the Gene Expression Omnibus (GEO), where it will be associated with an accession number. All data pertaining to modeling will be available through http://3DGenomes.org/datasets.

## Materials and Methods

### Mining for Oligopaint targets

Akin to a growing category of oligo-based FISH probes (e.g., ref. (52–56)), Oligopaint FISH probes consist of oligos that have been computationally designed to target specific sequences of the genome ((19); see the Oligopaints website (https://oligopaints.hms.harvard.edu/) for preselected whole genome datasets of Oligopaint oligo targets for the human, mouse, zebrafish, *Arabidopsis, Drosophila*, and *C. elegans* genomes). We identified genomic sequences that would serve as good targets for Oligopaint oligos by applying Oligominer to a repeat-masked human hg19 assembly (57). Specifically, we used the ‘balance’ settings of 35–41 nts of genome homology, 42–47°C for T_*M*_, a simulated hybridization temperature of 42°C, and ‘LDA mode’ filtering. Candidate probes were further processed by the ‘kmerFilter’ script that calls the algorithm Jellyfish (58) to eliminate probes containing 18mers that occur in >5 times in hg19.

### Mining the genome for homolog-specific Oligopaint (HOP) targets on chromosome 19 of PGP1

As the genome of PGP1 has been fully sequenced and phased (21, 59) (Kun Zhang and Daniel Jacobsen, unpublished; Bing Ren and Anthony Schmitt, unpublished), we were also able to design HOP probes for the homologs of chromosome 19; also see paper by O’Keefe, *et al*. (1996, ref.(60)). First, genomic sequences that would be good targets for HOPs (12) were discovered using the approach described above, except that the input hg19 sequence was not repeat-masked. These probe sequences were then intersected with the locations of phased heterozygous SNVs for PGP1 according to Lo, *et al*. (2013, ref.(59)) using the ‘intersectBed’ utility from BEDTools (61) and then processed with in-house software to generate two sets of haplotype-specific HOP probes in which each probe oligo targets ≥1 heterozygous SNV.

In order to ascertain which of the two HOP probes corresponds to maternal variants and which corresponds to paternal variants, we took advantage of the full genome sequence of PGP95 (courtesy of Rigel Chan and Elaine Lim), with whom PGP1 shares his maternal parent but not his paternal parent, and presumed that the homolog of chromosome 19 that carries one (or more) long blocks of variants in common with PGP95 would be the one that was maternally derived (Fig. 1B). First, heterozygous variants that identify the two PGP1 haplotypes were matched to variants from the full genome sequence of PGP95, and any that were homozygous in the PGP95 genome were discarded. All remaining PGP95 variants were then assigned as matching one of the PGP1 haplotypes (arbitrarily designated as H0 or H1) or neither, permitting us to identify blocks of contiguous matches to H0 or H1. The H1 haplotype included significantly more of the longest blocks shared between PGP1 and PGP95, thus identifying the H1 haplotype as maternally derived (Fig. S11). Then, as many long blocks corresponding to H1 were discovered to occupy a segment of PGP1 chromosome 19 at coordinates chr19:48,932,903-59,087,560, we designated the chromosome 19 homolog that carried these blocks the maternal chromosome. Accordingly, we designated the HOP probe corresponding to H1 as HOP-Maternal (HOP-M) and that corresponding to H0 as HOP-Paternal (HOP-P). The probes are efficient. Up to 96.5% of nuclei producing two fluorescent foci when imaged with HOP-M or HOP-P showed one of the signals to be stronger than the other, with the focus labeled more strongly with one probe being the more weakly labeled focus when imaged with the other probe; dye-swap experiments suggested that the difference between ratios for HOP-M and HOP-P is due to dye chemistries (Fig. S10B, C). Importantly, the corresponding ratio of signals obtained with probes targeting only the interstitial regions lying between SNVs approached 1.

### Multiplexing of the Oligopaint library and design of streets

We designed our Oligopaint libraries based on Hi-C maps that were available at the time our study was initiated. In particular, Geoffrey Fudenberg used IMR-90 Hi-C contact frequencies from Rao, *et al*., (2014, ref.(8)) to identify compartmental intervals and contact domains in the 8.16 Mb region extending from chr19:7,400,000 (19p13.2) to chr19:15,560,000 (19p13.12), where the calling of compartmental intervals relied on ICE (62) and a 3-state Gaussian Hidden Markov Model (HMM) implemented in scikit-learn (63), while contact domain calling relied on (64). Having segmented the 8.16 Mb region, we then incorporated Mainstreets and Backstreets into our library such that they would permit individual genomic features to be separately imaged. In particular, streets were generated using an in-house MATLAB (MathWorks, Natick, MA) script, ‘MakingStreets’, available via GitHub (https://github.com/gnir/OligoLego). Street sequences were vetted for predicted performance when serving as primers in PCR reactions using Primer3Plus (65) with default settings, except that the melting temperature was set to be 57-59°C, and the GC content was set to be 40-60% (65). Street sequences were also checked for pairwise orthogonality using NUPACK (66) to estimate equilibrium hybridization yields between candidate streets and other potential target sites in 390 mM Na+ and 50% formamide at room temperature (RT). Toehold sequences were also validated using NUPACK. Candidate streets passing all upstream checks were then examined for orthogonality to the human genome by using bowtie2 (67) to filter sequences aligning at least once to hg38 with ‘—very-sensitive-local’ alignment mode. Mainstreet and Backstreet sequences were appended to Oligopaint oligos using an in-house MATLAB code available via GitHub (https://github.com/gnir/OligoLego).

### Multiplexing Oligopaint libraries to walk along multiple chromosomes simultaneously

Here, we designed Mainstreet barcodes so that a) they could be shared across multiple chromosomes, such that one round of imaging could target anywhere from one up to the maximum number of chromosomes to be imaged, and b) imaging different chromosomes would be initiated in different rounds, thus permitting each chromosome to be identified by the round in which it first appears. For example, in an imaging scheme that introduces a new chromosome in every round, one could use the following scheme, wherein the walk along Chromosomes 1, 2, 3, 4, 5, and N are initiated in Rounds 1, 2, 3, 4, 5, and N, respectively.

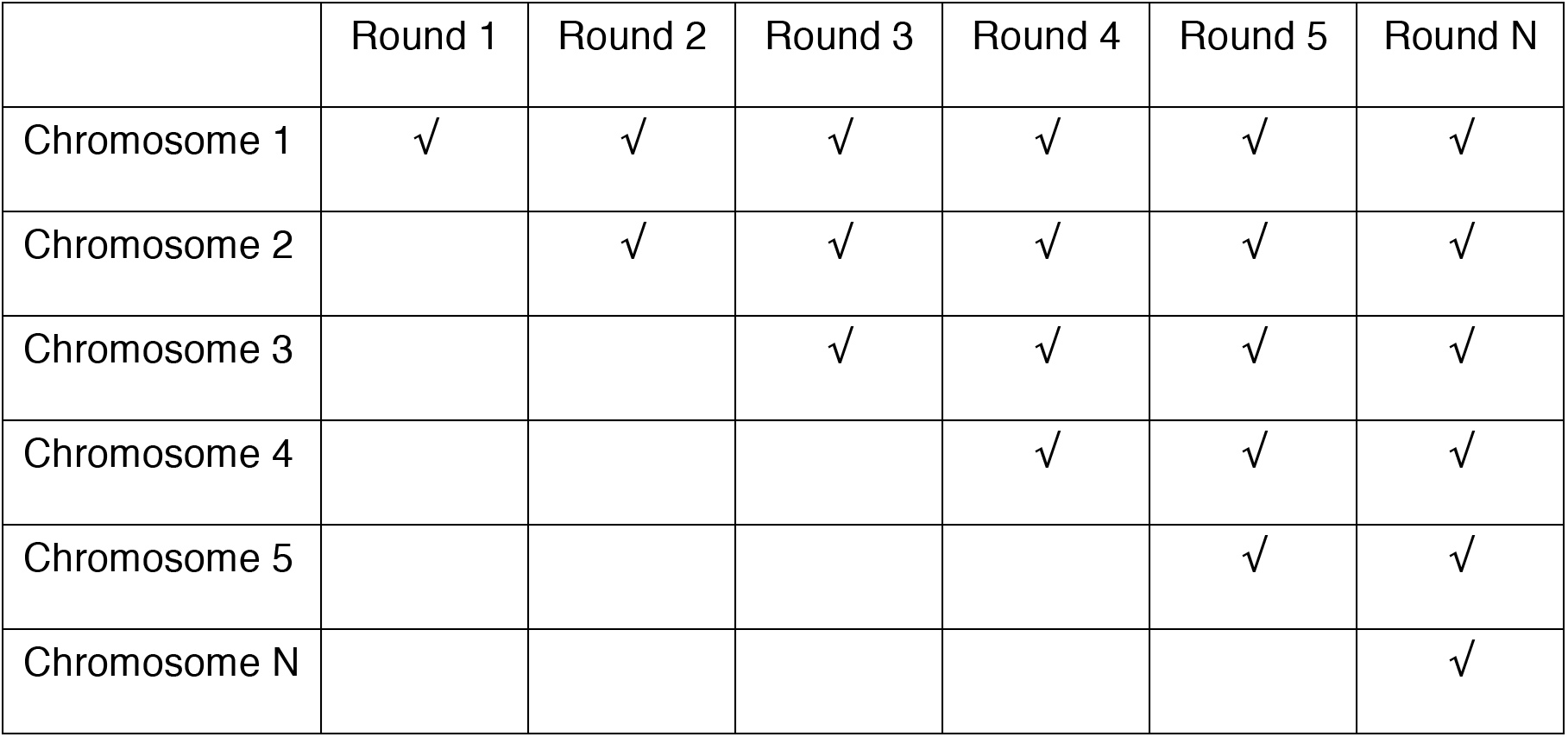

### Oligopaint probe synthesis

Oligopaint libraries were purchased from CustomArray (Bothell, WA) and amplified using *in vitro* transcription as previously described (11, 14, 20). Briefly, for each sub-pool to be amplified, we first optimized the template and primer concentrations using real-time PCR. We next performed large-scale limited-cycle PCR, and then used the product as template for *in vitro* transcription by T7 RNA polymerase (NEB, E2040S). The resulting ssRNA was then reverse transcribed (EP0752, ThermoFisher Scientific). RNA was digested by adding a mixture of 0.5 M NaOH and 0.25 M EDTA of equal volume to the reverse transcription reaction at 95° C for 10 minutes. Oligopaint ssDNA oligo probes were purified using the DNA Clean & Concentrator–100 kit (Zymo Research, Irvine, CA) paired with an EZ-Vac Vacuum Manifold (Zymo Research).

### Culturing of cells

Human primary skin fibroblasts of donor PGP1 (GM23248, Coriell Institute; (68)) were grown at 37°C + 5% CO_2_ in serum-supplemented (10% v/v) Dulbecco’s Modified Eagle Medium (DMEM) (serum Gibco 10437; media Gibco 10564). The cells were also supplemented with 1% (v/v) MEM Non-Essential Amino Acids Solution (Gibco 11140050). Penicillin and streptomycin (Gibco 15070) were also added to the cell culture media to final concentrations of 50 units/ml and 50 μg/ml, respectively.

### Sample preparation

40 mm coverslips *#*1.5 (Bioptechs, Butler, PA) were cleaned with a bath sonicator and stored in a 75% (v/v) ethanol solution at room temperature until used. In order to prepare PGP1 samples for imaging, the pre-cleaned coverslips were first allowed to fully dry in a tissue culture hood and then placed in sterile 150 mm tissue culture dishes (Falcon 353025). PGP1 cells were then deposited at ~15 % confluency onto the 150 mm dishes and allowed to grow in a mammalian tissue culture incubator until ~85% confluent (~5 days). At this point, the PGP1 cells were rinsed with 1x PBS, fixed using 4% (w/v) paraformaldehyde in 1x PBS for 10 minutes, rinsed with 1x PBS, permeabilized with 0.5% (v/v) Triton X-100 in 1x PBS for 10 minutes, rinsed with 1x PBS, and then stored in a cold room for up to 3 weeks.

### 3D DNA FISH

3D DNA FISH (69) was performed on fixed and permeabilized coverslips essentially as previously described (12, 19, 70). Briefly, coverglass samples were placed in small glass-bottom tissue culture dishes (MatTek P50G-1.5-30-F) and incubated with PBST (1x PBS + 0.1% v/v Tween-20) for 2 minutes. Samples were then incubated once with 1N HCL for 5 minutes, twice with 2x SSCT (2x SSC + 0.1% v/v Tween-20) for 1 minute, once with 2x SSCT + 50% (v/v) formamide for 2 minutes, and then once for 20 minutes with a pre-heated solution of 2x SSCT + 50% formamide at 60°C. Coverslips were then air-dried and incubated with a hybridization cocktail consisting of 2x SSCT, 50% formamide, 10% (w/v) dextran sulfate, 0.4 μg/μL of RNase A (ThermoFisher EN0531), and Oligopaint probe whose concentration was adjusted such that the final amount of probe added was equivalent to ~1.4 μM per every 1 Mb targeted (e.g. ~2.8 μM for probe targeting 2 Mb). In cases where the parental origin of the homolog was to be determined, HOPs probes were added at a concentration of 2.85 μM each. The hybridization reaction was then sealed beneath a 22 x 30 mm, *#*1.5 coverslip (VWR, Randor, PA) using rubber cement (Elmer’s, Westerville, OH), which was allowed to dry at 37°C for 7 minutes. Samples were then denaturated on top of a water-immersed heat at 80°C for 3 minutes. Coverslips were then allowed to hybridize for 2–3 nights at 47°C in a humidified chamber. Following hybridization, coverslips were washed 4 times with 2x SSCT at 60°C for 5 minutes, then twice with 2x SSCT at RT. Coverslips were then rinsed with 1x PBS and allowed to air dry. For the purposes of drift correction, fiducial markers (gold nano-urchins, GNUs; d=90 nm, 630 nm abs max) (Cytodiagnostics, Ontario, Canada) were sonicated for 10 minutes and then diluted 1:3 in PBS, pipetted onto the sample coverslips, and sandwiched with a blank, i.e. nontreated, 40 mm coverslip. GNU-coverslip sandwiches were centrifuged at 500 x *g* for 3 minutes. The sample coverslips were then washed for 2 minutes with 1x PBS.

### HOPs imaging and processing

Images were obtained using the widefield mode on the Bruker Vutara 352 (Bruker Nano Inc.) microscope (described in the section in Methods on OligoSTORM imaging). *SlowFade* antifade reagent containing DAPI dye (ThermoFIsher Scientific) was used as the imaging buffer. Images were acquired in the Z-dimension, approximating 2-3 um across 4 different channels, with the following dyes: DAPI, Alexa Fluor 647, Alexa Fluor 488, and ATTO 561 with 405, 488, 561, and 639 nm lasers (Coherent), the emission filters (custom-made, Semrock) 465/30, 515/40, 600/50, respectively, and detected via an sCMOS camera (Hamamatsu Orca-flash 4.0 v3) with 100 ms exposure time. These four channels corresponded to nuclear staining, interstitial signals, and the two HOP probes, respectively. Images were analyzed using an inhouse MATLAB (Mathworks) script, which was written to detect and then derive the intensity ratio of the foci in each channel. Detection of signal from foci was validated by eye.

### OligoSTORM imaging

OligoSTORM imaging was performed on a Bruker Vutara 352 commercial 3D biplane single molecule localization microscope (71–73), equipped with a 60x water objective (Olympus) with a numerical aperture (NA) of 1.2. For illuminating Oligopaint oligos with Alexa Fluor 647, we used a 639-nm Coherent Genesis laser with ~ 25 kW/cm^2^ at the back aperture of the objective, and for illuminating with Alexa Fluor 405, we used a 405-nm Coherent Obis laser for photoactivation with up to ~0.05 kW/cm^2^ at the back aperture (emission filters were described in the Methods section describing HOPs imaging and processing). Fluorescent detection was captured on an sCMOS camera (Hamamatsu Orca-flash 4.0 v2) using a 10 ms exposure time. At each imaging session, we imaged 5-10 nuclei at different X,Y stage locations, while taking up to 4 μm z-scan using 100 nm Z-steps at each location, with up to 21 loci per nucleus, collecting 42,000-150,000 frames for each locus. Image data was collected by a Bruker-provided network attached storage (NAS) system.

### OligoDNA-PAINT

DNA-PAINT imaging was performed on a custom optical set-up based on a Nikon Ti Eclipse microscope featuring a Perfect Focus System and a custom-built TIRF illuminator. DNA-PAINT was performed using Highly Inclined and Laminated Optical sheet illumination (HILO; (74)) with 10% of a 1000-mW, 532-nm laser (MPB Communications) using a CFI Apo TIRF 100× oil (N.A. 1.49) objective at an effective power density of ~2 kW/cm^2^. The 532-nm laser excitation light was passed through a quarter-wave plate (Thorlabs, WPQ05M-532), placed at 45° to the polarization axis and directed to the objective through an excitation filter (Chroma ZET532/10x) via a long-pass dichroic mirror (Chroma ZT532RDC_UF2). Emission light was spectrally filtered (Chroma ET542LP and Chroma ET550 LP), directed into a 4f adaptive optics system containing a deformable mirror (MicAO 3DSR, Imagine Optic) and imaged on an sCMOS camera (Andor Zyla 4.2+) with 6.5-μm pixels using rolling shutter readout at a bandwidth of 200 MHz at 16 bit, resulting in an effective pixel size of 65 nm. In order to estimate the axial positions of single molecule emitters, optical astigmatism was applied using the deformable mirror to cause an asymmetric distortion in the observed point spread functions (75, 76). A total of 22,500–45,000 250-ms frames were acquired for each image by using 0.5–1 nM of Cy3B-labeled 10-mer oligos in 1× PBS solution + 500 mM NaCl, 10 nM PCD, 2.5 mM PCA, and 2 mM Trolox. Gold nanoparticles (40 nm; no. 753637; Sigma-Aldrich) were used as fiducial markers to facilitate drift correction. The lateral positions of single-molecule localization events were first determined using Picasso (77) by applying the least-square fitting algorithm, and the axial positions were then also determined using Picasso by fitting to a pre-established calibration curve (75, 76). Global lateral drift-independent localization precision was estimated by Nearest Neighbor based Analysis (NeNA) (78). Global axial localization precision was estimated by empirical analysis of sub-diffraction sized fiducial markers after fitting to an empirically derived calibration curve with a mean precision of < 20 nm.

### OligoDNA-PAINT image reconstruction

Two-dimensional DNA-PAINT images were rendered using Picasso, with individual localization events being represented as single point sources blurred by an isotropic twodimensional Gaussian function whose “sigma” parameter reflects the average X-Y NeNA localization precision of the localizations occurring in the presented field of view and colored according to inferred axial position (78). Multicolor three-dimensional DNA-PAINT images were rendered using ViSP (79), with individual localization events being represented as single point sources blurred by an anisotropic three-dimensional Gaussian function whose X-Y “sigma” reflects the average NeNA localization precision of the localizations occurring in the presented field of view and Z “sigma” was informed by the precision of the calibration curve used for axial fitting as determined by Picasso.

### Automated microfluidics

We exploited a fluidics system as previously described (11, 23). Briefly, Bioptech’s FCS2 flow chamber (v ~ 100 μL) was integrated with a peristaltic pump (Gilson minipuls3) and 3 valves (Hamilton HVXM 8-5), with the resulting dead volume in our system being ~700 μL. Integration of the fluidics system and the Bruker Vutara 352 microscope was programmed into SRX, the commercially available microscopy software package that controls the Bruker Vutara 352.

### Sequential hybridization for multiple rounds of OligoSTORM

Secondary hybridizations were performed using 50 μM of secondary probe in 2x SSC +30% (v/v) formamide, and allowed to hybridize for 30 minutes at RT. Following hybridization, a wash solution (40% v/v formamide + 2x SSC) was introduced to the flow cell and incubated for 3 minutes without flow. Wash solution was then replaced with imaging buffer consisting of 10% (w/v) glucose, 2x SSC, 50 mM Tris, 1% (v/v) β-mercaptoethanol, and 2% (v/v) of a GLOX stock solution consisting of 75 mg/mL glucose oxidase (Sigma G2133-250KU), 7.5 mg/mL catalase (Sigma C40-500MG), 30 mM Tris, and 30 mM NaCl. A thin layer of mineral oil (Sigma M5904) was added at the top to prevent oxygen from penetrating the imaging buffer. Imaging was initiated after the imaging buffer was allowed to incubate for 2 minutes without flow.

### OligoSTORM data processing

During localization analysis, the precision of each localization event was determined by calculating the Cramér-Rao Lower Bound, namely the inverse of the Fisher information of the measured point-spread function (80, 81). The Cramér-Rao Lower Bound is the lower bound of the variance of the estimation process, which is used to calculate the localization precision of each event.

For isolating localizations comprising structures of interest during each round of imaging (fluidic cycle), we first removed localizations of lesser quality according to the following criteria: initial filtering of the localization data consisted of applying a geometric mean goodness-of-fit filter calculated from three separate metrics from the localization data. The individual metrics consisted of calculating the ratio of the photons assigned to the Point Spread Function (PSF) in the localization routine to the total number of photons in the camera cutout around a localization event (first metric), total photons in the cutout around a localization event (second metric), and the offset in Z between the calculated localization position of the fluorophore and the physical position of the objective piezo (third metric). The goodness-of-fit metric of each localization event is calculated by sorting each metric through a self-normalization process (with a value of 0 and 1 being the lowest and highest, respectively, quality value to a given metric), multiplying the values of the three metrics together, and taking the cube-root. Localizations with a score closer to 1 are considered of higher quality as compared to localizations with a score closer to 0. An initial thresholding was set such that only localizations with a rank of 0.8 or higher were accepted for statistical analysis. Also, candidate localization events whose peak intensity varied no more than 2 pixels (pixel size ~100 nm) in up to 7 frames in a row were considered as one. Finally, we filtered out all localizations with an axial precision worse than 100 nm. We found that the number of localizations increases with genomic size (Fig. S12). Following localization filtering, we performed drift correction using a center-of-mass function between subsets of fiducial markers across the data recording (82). Finally, clusters of localizations were segmented using DBSCAN (34).

When determining the spatial overlap between two clusters, we used Bruker’s SRX software to generate surfaces of the calculated clusters based upon alpha shapes, using an alpha radius of 150 nm, which reflects the maximum particle distance used for the DBSCAN analysis for CS1, CS7, and the merged four contact domains that were imaged in rounds 10-13 (Fig. 2A, H). Average spatial overlap between two clusters was expressed as the average of the fraction of the first cluster overlapping the second and the fraction of the second cluster overlapping the first.

For comparison, we determined the spatial overlap of a cluster that was randomly divided into two clusters. In particular, we chose two nuclei as an example and divided the localizations obtained in rounds 1 and 21 of imaging into two, generating 8 clusters per diploid nucleus. We noticed that for the nucleus with the higher number of localizations (~6,500 localizations/cluster) the average spatial overlap was 90-92%, while for the nucleus with lower number of localizations (~3,380 localizations/cluster) the average spatial overlap was 85-89%. This implies that the spatial overlap between rounds is not expected to be more than 90% and is dependent upon the number of localizations.

### Selection of nuclei for analysis

For OligoSTORM analysis, we chose only diploid nuclei for which the homologs were not too close to be distinguished with a single dye (as we used only Alexa647N for OligoSTORM), did not show high levels of anisotropy in Z, and also did not show evidence of aneuploidy. Note, PGP1 cells were confirmed to be karyotypically normal (XY; n=20; Cell Line Genetics, Madison, WI), and cell sorting experiments show that 72-86% of PGP1f cells are in G1.

### Generating OligoSTORM density maps

DBSCAN-extracted localizations for each chromosomal segment were convolved with a Gaussian kernel to match the mean precision (37) and to reduce the anisotropy of the image acquisition. The density map of each homolog compartmental segment was represented by intensities at points *i* (ρ_i_) on a cubic grid with fixed voxel size. For each localization, the density value of the nearest voxel was increased by a factor of 1. The obtained grid was convoluted with a Gaussian function using TEMPy (37), such that the density ρ_i_ was defined as:

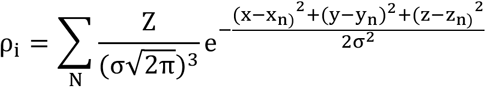

where *x, y* and *z*, and *x_n_, y_n_, z_n_*, are the Cartesian coordinates of grid point *i*, and localization *n* respectively. *N* is the total number of localizations, *z* is the number of localizations per grid point, and *σ* is set to be proportional to the mean precision. The iso-contour threshold for each map was set to be one sigma after shifting of the background peak to zero using TEMPy (37). The density map of the entire walk was also generated using the same approach.

### Analysis of OligoSTORM density maps

A total of five structural measures were obtained from each OligoSTORM density map obtained for each chromosomal segment. First, the distance score (DS) between two chromosomal segments was calculated as the spatial distance between the center of mass of each OligoSTORM density map. Next, the DS was corrected for linear genomic distance using a power-law fit. Finally, the DS Z-score was calculated for each data point using the power-law function as expected value. Second, the entanglement score (ES) between two chromosomal segments was calculated as the fraction of overlapping voxels within the optimal iso-contour threshold with respect to the smaller of the two density maps as implemented in TEMPy (37, 83). As before, the ES was corrected for linear genomic distance using a power-law fit obtained by the ES Z-score. Third, the surface area of each OligoSTORM density map was calculated by summing over the area of the triangles on the surface on the convex hull using the “measure area” option in Chimera (50). Next, the surface area was corrected for genomic size (Fig. S6A) and transformed into a surface area Z-score. Fourth, volume was calculated within the optimal iso-contour threshold using the “measure volume” option in Chimera (50). As before, the volume was corrected for genomic size (Fig. S6B) and transformed into a volume Z-score. Fifth, a sphericity score (*Ψ*), which measures how closely the shape of an OligoSTORM density map approaches that of a mathematically perfect sphere, was calculated as:

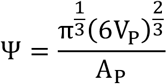

where *V_P_* and *A_P_* are the volume and surface area of the OligoSTORM density Map *P*. To take into account the bias due to genomic size, sphericity was corrected for genomic size (Fig. S6C) and transformed into a sphericity Z-score.

For each homolog, a feature vector was created, which is a binary 1×9 vector encoding the cluster types of each chromosomal segment in the region. Next, a per-nucleus profile of the compartment state was made for comparing the feature vectors of each of the homologs in a single nucleus. The compartment state profile is represented by a 1584:3429 vector whose values are determined by a comparison score set as: (i) 1, both compartment segments belong to cluster 1, (ii) 0, the compartment segments belong to different clusters, or (iii) −1, both compartment segments belong to cluster 2. The resulting matrix of 19 compartment state profiles was hierarchically clustered, resulting in five clusters.

The resulting matrix of structural features for the 342 chromosomal segments (that is, 9 segments in 2 homologs in 19 cells) was analyzed using the PCA implemented in the Python library Scikit-learn. The clustering resulted best in two major clusters that were named cluster 1 and cluster 2 (Fig. 3C, S7A).

### ENCODE PGP1 data

Data available for RNA-seq, DNase-seq and five chromatin marks (H3K4me1, H3K27ac, H3K4me3, H3K27me3, H3K9me3) were downloaded from the ENCODE project (46). Mapped reads were obtained from samples ENCFF043JNP and ENCFF280JPQ for H3K4me1, ENCFF612WGP and ENCFF345SWP for H3K27ac, ENCFF397XVM and ENCFF179TEP for H3K4me3, ENCFF378EQH for RNA-seq, ENCFF523JNF and ENCFF050OVJ for DNase-seq, ENCFF800CCA and ENCFF876WHO for H3K27me3, and ENCFF644QGC for H3K9me3 (https://www.encodeproject.org). Next, coverage of the chr19:7 (19p13.2) to chr19:15.5 (19p13.12) visualized region was calculated using the SamTools bedcov command (84) at 10 kb resolution. All datasets, with the exception of RNA-seq and H3K9me3 ChIP-seq, had two replicates which were highly correlated (r > 0.99) and, in such cases, the average coverage per each bin was used. The final number of mapped reads for each dataset was 60,550,146 for H3K4m1, 52,286,921 for H3K27ac, 62,084,095 for H3K4m3, 200,776,708 for RNA-seq, 622,389,260 for DNAse, and 65,357,125 for H3K27me3, and 26,947,714 for H3K9me3.

### Homolog ellipticity

The Cartesian coordinates of the location of each homolog in the cell population were fitted to an ellipsoid (85), and the ellipticity score for any 8.16 Mb region was then calculated to be the ratio between the two largest principal axes of the ellipsoid. When comparing homologous chromosomal regions in a single nucleus, we determined the ratio between the more elliptical homolog and the less elliptical one, calling that measure the ellipticity ratio. An ellipticity ratio equal to 1 indicates that the two homologs of a single nucleus had the same ellipticity score. For the evaluation of significance, a random ensemble of elongation ratios was constructed by calculating the ratio between two homologs randomly selected from the analyzed cell population.

### Modeling of genomes

#### Integrative 3D modeling with TADbit

The chromosome 19 normalized Hi-C interaction matrices of each chromosomal segment and the entire region (chr19: 7400000:15560000) were used for modeling at a resolution of 10 kb as previously described (41, 42). The selected resolution was the highest possible with an MMP score predicted to result in reliable 3D models ((86) and Table S7). For each chromosomal segment, two independent ensembles of models were generated: (i) Isolated chromosomal segments and (ii) the chromosomal segments extracted from models of the entire region. Briefly, TADbit automatically generates 3D models (42) using a restraint-based modeling approach, where the experimental frequencies of interaction are transformed into a set of spatial restraints (41). For each chromosomal segment, a total of 5,000 models were generated, with only the best 1,000 models (that is, those that best satisfy the input restraints) used for further analysis. For each of the modeled chromosomal segments, the contact map obtained from the final 1,000 models resulted in a good correlation with the input Hi-C interaction matrix (Table S7), which is indicative of good model accuracy (86).

#### Rigid fitting

The resulting 3D models for each chromosomal segment ensemble were rigidly fitted in the corresponding OligoSTORM density maps using the ‘Fit_in_map’ global rigid-body fitting protocol in Chimera (50). A total of 100 random placements of each chromosomal segment were generated by randomly rotating and translating the fit within the reference OligoSTORM density map, followed by a step of local optimization using steepest ascent. Next, two scores were computed for each chromosomal segment fit: (i) the cross-correlation coefficient (CCC) between the OligoSTORM density map and the chromosomal segment fit and (ii) the connectivity score (ConS) of the chromosomal segment fit with that of its sequentially neighboring chromosomal segment. The CCC score is expressed by the following formula (37, 87):

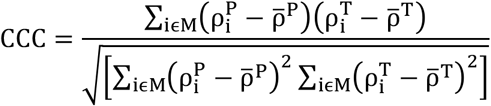

where *M* is all the voxels in the density grid of the map; *P* and *T* are the chromosomal segment fit map and OligoSTORM density map, respectively; 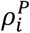 and 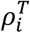 are the intensity of the density maps at voxel *i*; and 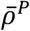 and 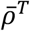 are the respective mean intensity values. The CCC ranges from 0 (no match) to 1 (perfect match). The ConS, which ranges from 0 to 1 was defined as:

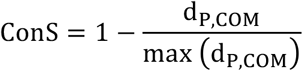

where *d_P,COM_* is the distance between the starting or end bin of the fit with the center of mass of the surface interface with its neighboring chromosomal segment, and *max*(*d_P,COM_*) is the maximal *d_P,COM_* in the ensemble of fits. The surface interface is defined in a multistep process: (i) identification of the surface area of the chromosomal segment in analysis that lies within a cut-off distance of 50 nm from its neighbor chromosomal segment and calculated by the “measure contactArea” option in Chimera (50), and (ii) selection of the density points of the chromosomal segment density map that are bounded by the identified surface area by using the “mask” command in Chimera (50). Finally, a combined score (Fig. S13) integrating both measures was used for calculating the goodness of fit:

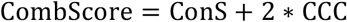

For each chromosomal segment, the top-scoring fits (95^th^ percentile) were chosen for further refinement.

#### Flexible refinement

To further improve the agreement between the fit and the OligoSTORM density maps, a flexible fitting method was implemented similarly to the protocols for conformational refinement of protein guided by cryoEM density (43). The protocol is a heuristic optimization that relies on simulated annealing molecular dynamics applied to a series of subdivisions of the fit into progressively smaller rigid bodies (*b_l_*). A rigid bod *b_l_*y could be any set of particles in the fit (including loops, a sub-domain, a contact domain, compartment segment or larger region). A set of rigid bodies (*B*) was defined for each fit using TADbit’s TAD caller and selecting borders with strength higher than 5. Finally, the exact rigid bodies were manually adjusted by visual inspection of the rigid fitted 3D starting model (Table S8). During the refinement, the coordinates of each *b_l_* ∈ *B* were displaced in the direction that maximized their cross-correlation with the OligoSTORM density map and avoided the violation of the imposed restraints from the population Hi-C interaction matrix. The final scoring function was defined as:

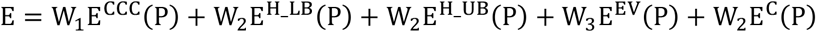

where *E^CCC^*(*P*) quantified the fit of the probe map (map generated from the 3D fitted model) (*ρ^P^*) and the target OligoSTORM density map (*ρ^T^*), and is defined as the negative sum of crosscorrelation coefficients between the OligoSTORM density map and the rigid bodies *b_l_*; *E^H_LB^*(*P*) quantified the lower-bound harmonic restraints which prevents two particles from getting closer than a given equilibrium distance; *E^H_UB^*(*P*) quantified the upper-bound harmonic restraints, which forces two particles to be closer than a given equilibrium distance; *E^EV^*(*P*) is an hard sphere excluded volume term (lower bound harmonic oscillator dropping at zero at 2r); and *E^C^*(*P*) is a harmonic bond that connects adjacent particles at a given equilibrium distance. The weights *W*_1_, *W*_2_, and *W*_3_ determine the relative importance of the corresponding terms and were set as default, which proved optimal for fitting of sub-tomogram averaging (88). The optimization protocol to identify the best fit consisted of simulated annealing rigid-body molecular dynamics followed by a final a conjugate-gradient minimization. A maximum of 50 cycles of simulated annealing molecular dynamics steps were performed, in each cycle the system was heated from 0K to 1000K for a maximum of 100 steps and cooled back to 0K for a maximum of 200 steps. At each step, the optimization was terminated if the change in CCC was < 0.001. Finally, a minimization step was performed comprising 200 conjugate-gradients steps with weights *W*_1_, *W*_2_ and *W*_3_ set to 1 and 200 conjugate-gradients steps with *W*_1_set to 0 and *W*_2_ and *W*_3_set to 1. The goodness-of-fit was assessed using the CCC score and the clash score (CLS). The latter defined as the number of serious clashes (that is, distance *d*<2r) per 1000 beads. The refined model was chosen as the one that maximizes the CCC and minimizes the clash score (Table S8).

### PGP1F Hi-C

PGP1f cells were cultured as described in the ‘Cell Culture’ section of Methods and pelleted at low speed centrifugation (185 x g) to maintain cellular and nuclear integrity. We generated 2 *in situ* Hi-C libraries with three million cells each, utilizing the MboI restriction enzyme (NEB, R0147M), following the protocol described in (8). Briefly, the Hi-C protocol entails crosslinking cells with 1 % formaldehyde (wt/vol), cell permeabilization with nuclei intact, DNA digestion with an appropriate 4-cutter restriction enzyme, and 5’ -overhang filling with the incorporation of a biotinylated nucleotide. The resulting blunt end fragments are ligated, and the DNA is then sheared. Biotinylated ligation junctions are captured with streptavidin beads, and the resulting fragments are analyzed with paired-end sequencing.

### PGP1f Hi-C data processing

The two Hi-C libraries were sequenced with 150 bp paired-end sequencing reads on an Illumina HiSeqX-10. The resulting sequencing data from both libraries (Table S9) was processed separately using Juicer (89) and mapped to the GRCh37/hg19 (hg19) human genome assembly (8, 89). To generate statistics and a Hi-C file for a combined map including both replicates (Table S5; Fig. S14), the mega.sh script was used (https://github.com/theaidenlab/juicer/wiki/Usage (89), with the following command: /gpfs0/juicer/scripts/mega.sh.

The combined Hi-C map generated a total of 691,620,560 reads, achieving a resolution of 5 kb, in alignment with Hi-C map resolution parameters described in (8). This combined map was used with ‘KR normalization’ (51, 89) for all subsequent analyses.

### Assignment of manually identified compartments as ‘A’ or ‘B’

To label the two manually identified compartments as ‘A’ or ‘B’, we began by partitioning chromosome 19 into 100kb loci. These loci were assigned to two compartments based on the intrachromosomal contact matrix for chromosome 19 using the eigenvector method (Fig. S8) as implemented in Juicer (3, 89). Specifically, we used the following command:

~~~
‘java -jar /gpfs0/juicer/scripts/juicer_tools.jar eigenvector KR inter.hic 19 -p BP 100000’.
~~~

Visual inspection of the Hi-C map for chromosome 19 confirmed that the eigenvector corresponded reliably to the plaid pattern in the data (3) (Fig. S8 and interactive Fig. S8 http://bit.lv/2zCG2X6, for interactive figure documentation see: https://igvteam.github.io/juicebox.js/).

Next, we labeled the two compartments as ‘A’ and ‘B’ on the basis of GC content, which is known to be enriched in the ‘A’ compartment and in open chromatin more generally (90). The GC content track for human hg19 chr19 was downloaded from the UCSC Genome Browser (http://hgdownload.cse.ucsc.edu/goldenpath/hg19/gc5Base/) and encodes the raw data for the gc5Base track on hg19 (site last updated on 24-Apr-2009 14:48). We binned the data at 100 kb resolution for chromosome 19, using the following awk command:

~~~
awk -v res=100000 ‘BEGIN{fname=“temp”}NF>2{split($2,c,”=“); if
(fname!=c[2]){print tot ≫ fname”.txt”; close(fname”.txt”); m=res;
tot=0} fname=c[2]; }NF==2 && $1>m{while ($1>m){print tot ≫
fname”.txt”; m=m+res; tot=0;}}$1<=m{tot+=$2; }END{print tot ≫
fname”.txt”; close(fname”.txt”);}’ my_file.wig
~~~

Finally, we calculated the Spearman rank correlation coefficient between the compartment eigenvector and the GC content track. We obtained a Spearman rho value of - 0.674928131634, which implies that the ‘A’ compartment corresponds to loci whose eigenvector entry is negative and the ‘B’ compartment corresponds to loci whose eigenvector is positive (Fig. S8, Interactive Figure http://bit.ly/2zCG2X6).

Accordingly, we labeled compartmental intervals 1, 3, 5, 7, and 9 (at coordinates 7 - 8.6 Mb, 9.3 – 9.5 Mb, 9.6 – 9.8 Mb, 9.9 – 14.6 Mb, and 15.2-15.5 Mb) as compartment ‘A’ and intervals 2, 4, 6, and 8 (with coordinates 8.7 – 9.3 Mb, 9.5 – 9.6 Mb, 9.8 – 9.9 Mb, and 14.6 – 15.2 Mb respectively), as compartment ‘B’. Note, compartmental intervals may differ from chromosomal segments by genomic coordinates and/or compartment classification.

## Supplementary notes

### Integrative Modeling Procedure

To trace the chromatin structure of the imaged loci, we implemented a fitting procedure that integrates super resolution data and Hi-C interaction maps. The starting point of the procedure is an OligoSTORM density map and an ensemble of 3D models generated based on the Hi-C PGP1f mega map using TADbit (41, 42). The approach proceeds in two main stages (Fig. S9A):

#### Step 1: Rigid Body Fitting

In this step, we search for the optimal position and orientation of each chromosomal segment 3D model in the ensemble in the respective OligoSTORM density map (Methods) using a six-dimensional search (three translational and three rotational degrees of freedom). During the rigid body fitting procedure, we evaluate the cross-correlation coefficient (CCC, Methods) between a simulated density map from the 3D model and the OligoSTORM density map. Two keys aspects will mainly influence this first step: (i) the number of distinct high-density feature points of the map and (ii) a good initial placement of the model in an OligoSTORM density map. The initial components are manually placed in the density map, and then, for each 3D model, 100 perturbations are performed and locally optimized within the density map by steepest ascent method (Methods). Using the rigid body fitting step, we can identify the approximate position and orientation of the chromosomal segment 3D model in the density map.

The goodness-of-fit in this step is assessed with a combined score (*`CombScore`*) of CCC with the density map and connectivity with the neighboring chromosomal segment (Methods and Fig. S10A, B). The top-percentile best-fitting models for each chromosomal segment OligoSTORM density map were selected for the second step of the integrative protocol (with the exception of CS5 in homolog 2, which was manually fitted due to a poor connectivity score with neighboring chromosomal segments).

#### Step 2: Flexible Fitting Refinement

In step 2, the conformation of the chromosomal segment was optimized simultaneously with its position and orientation in the OligoSTORM density map via flexible fitting refinement (43). Each model in the set of best-fitting models for each chromosomal segment was subdivided into a smaller rigid-body, in agreement with the manually curated contact domains from the Hi-C experiment (see Methods). Then, we optimized the 3D model by refining positions and orientations of its rigid bodies with a simulated annealing rigid-body molecular dynamics protocol for up to 50 cycles. The scoring function used for the dynamic simulation in this second step included the *E*^CCC^ term with the OligoSTORM density map and terms for the satisfaction of the spatial restraints (imposed as harmonic restraints) as calculated from the Hi-C experiment. The goodness-of-fit was assessed with the CCC and the clash (CLS) scores (Table S8). We chose these scores as it has been previously shown for proteins that the improvement in CCC can be associated with the goodness-of-fit of the final models (43) (however other goodness-of-fit measurement can be used (37, 91) and, furthermore, that the CLS scores are a good proxy for quantifying unfavorable steric overlaps. Generally, this stage of refinement is considered “finished” when the molecular dynamics simulation has converged as evaluated by the goodness-of-fit measures. The absence of such convergence could be related, for example, to a sub-optimal starting rigid-body conformation, an incorrect rigid-body assignment, and/or few high-density feature points in the map.

### Assessment of *in-situ* Hi-C data reproducibility

In order to assess reproducibility of our *in situ* Hi-C replicates, we calculated the Pearson’s r between Hi-C maps as a function of distance, as described in (8). Our data remained highly correlated at all resolutions tested (see Fig. S14B for correlation as a function of the distance). To do this, we utilized a MATLAB script, which can be found here https://www.dropbox.com/s/f8qezt6ny5v6yml/plt_correlation_distance.py?dl=0

### Supplementary figures

**Fig. S1.**
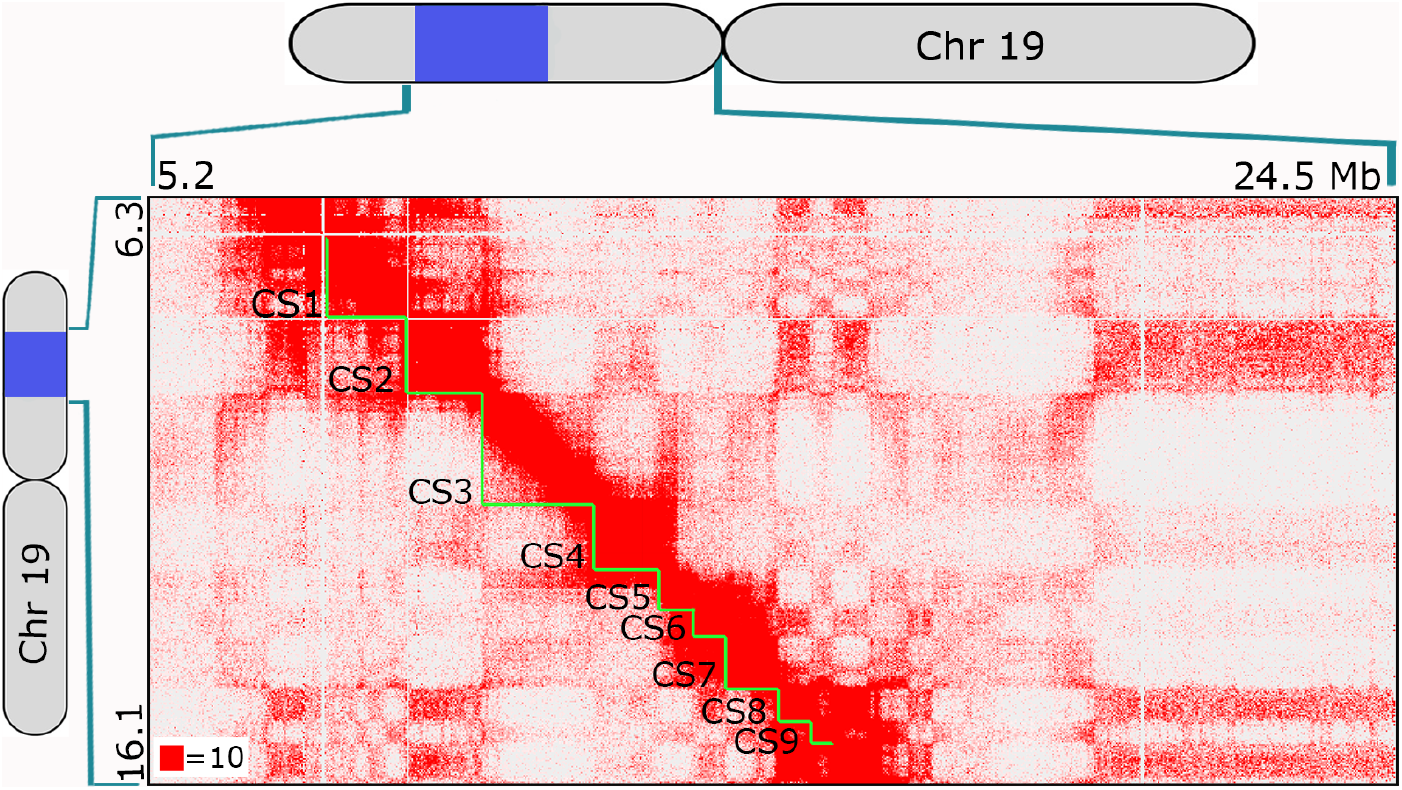
*In situ* Hi-C map of CS1 −9 in IMR-90. Hi-C map of region including CS1 −9, displayed at 25 kb resolution and normalized according to Knight & Ruiz (51). Contact count indicated by the color of each pixel, ranging from 0 (white) to 10 (red); OligoSTORM steps (CS1-9) delineated in green. Cartoons of chromosome 19 show extent of the 8.16 Mb imaged region (blue).

**Fig. S2.**
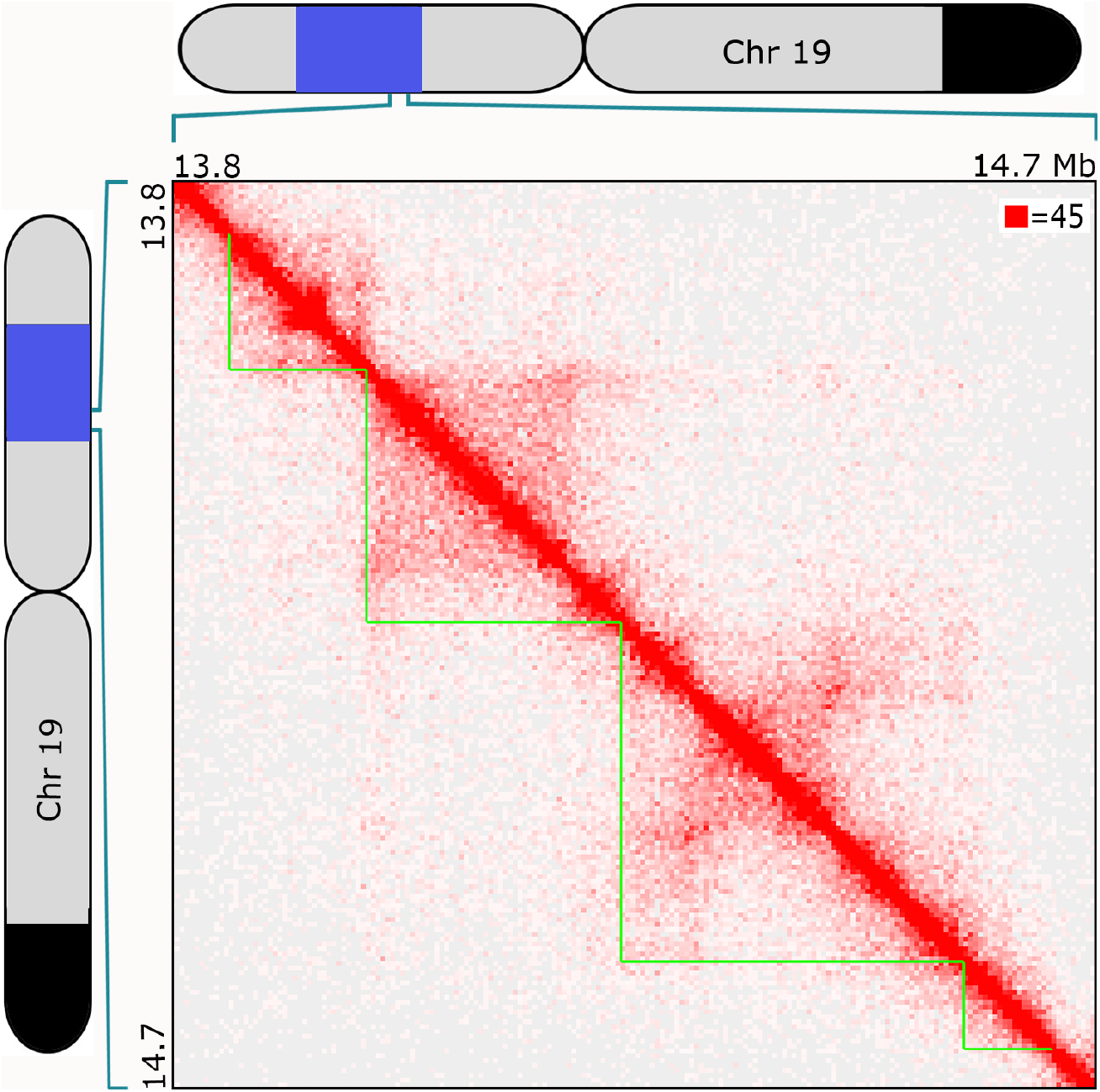
In situ Hi-C map of CS7 in PGP1f. Hi-C map of region including CS7, displayed at 5 kb resolution and normalized using Knight & Ruiz (51). Contact count indicated by the color of each pixel, ranging from 0 (white) to 45 (red); 4 subregions of CS7 delineated in green. Cartoons of chromosome 19 show extent of the 8.16 Mb imaged region (blue) and block of SNVs shared between PGP1 and PGP95 (black; as in Fig. 1B).

**Fig. S3.**
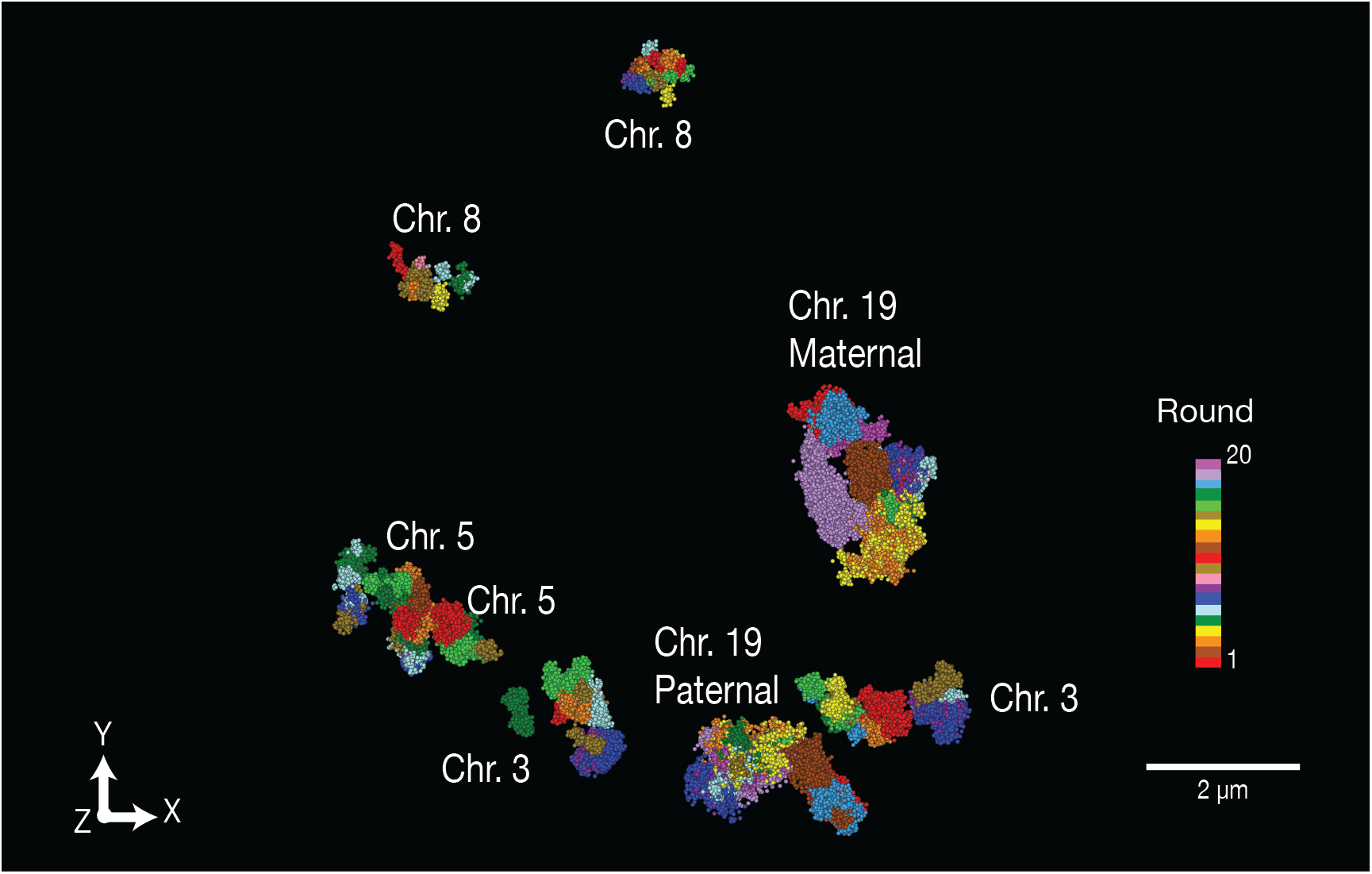
Walks along multiple chromosomes using OligoSTORM. Nucleus showing walks initiated in rounds 4, 5, and 6 on Chromosomes 8, 5, and 3 in step sizes of 100, 250, and 500 kb, respectively, while simultaneously walking along Chromosome 19, commencing in round 1 and in step sizes ranging from 3 to 1,800 kb. This walk encompassed a total of 23 cycles (see color bar) and, although it was not successful in all rounds on all chromosomes, it nevertheless illustrates the potential of this approach to increase throughput.

**Fig S4.**
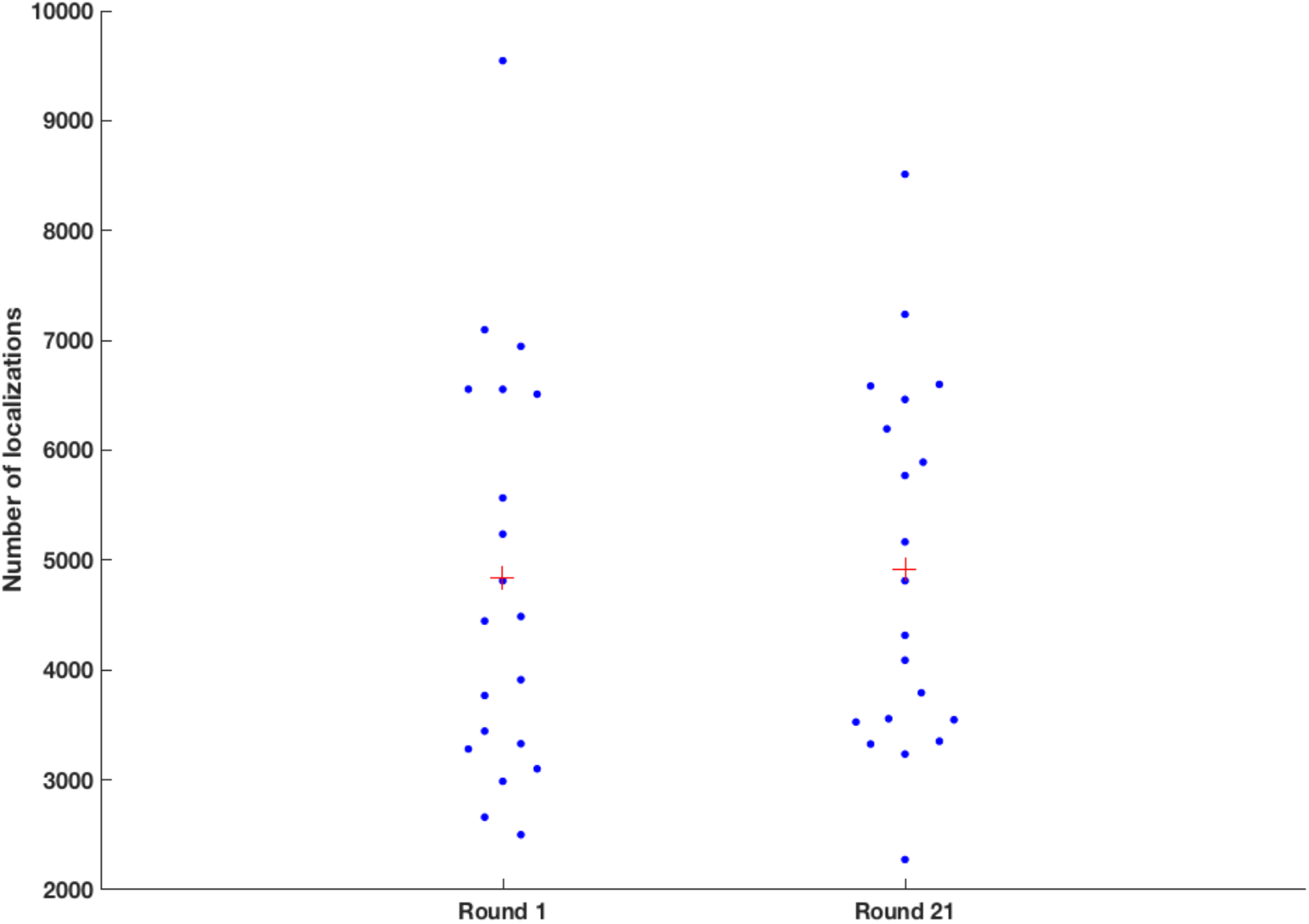
Comparison of number of localizations per cluster between fluidic cycles 1 and 21. After querying the change in fractional overlap for CS1 after 21 rounds of imaging with OligoSTORM (Fig. 2G), we assessed whether the number of localizations had changed using a Beeswarm plot (MATLAB script written by Jonas, MATLAB community) showing the number of localizations detected from 20 homologs (total of 10 nuclei, colored as blue dots). The mean number of localizations per cluster (red cross) is similar between rounds 1 and 21, implying no significant change (two-sample Kolmogorov-Smirnov test, p=0.9655).

**Fig. S5.**
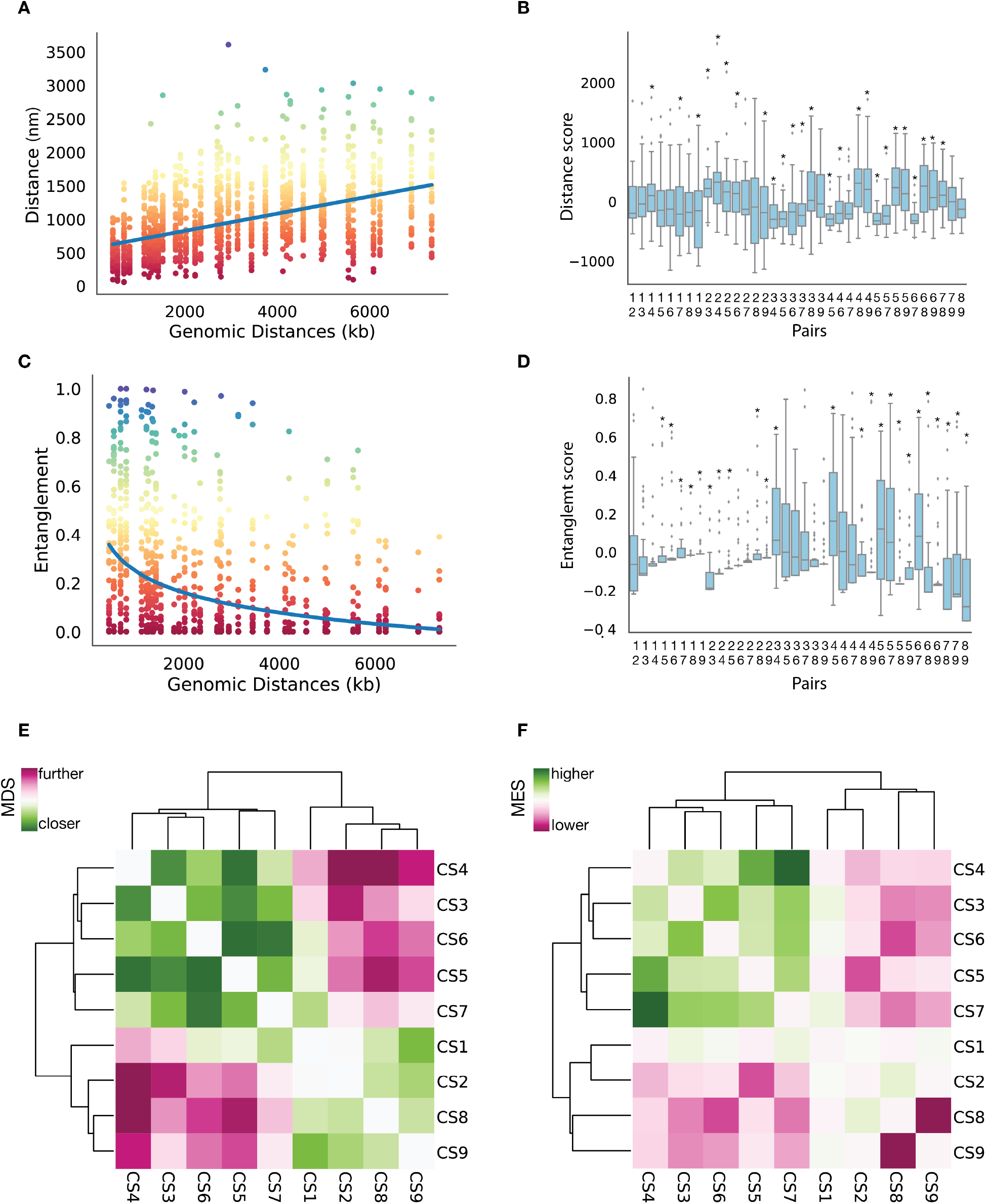
Variability of spatial distances and entanglement observed for chromosomal segments. (**A**) Distance versus genomic distance for the 342 analyzed chromosomal segments using a power-law function fitted to the data (blue line). Data are colored by their distance value (rainbow color palette). (**B**) Distribution and variation of the distance score (DS) of the two chromosomal segments being compared. (**C**) Entanglement versus genomic distance for the 342 analyzed chromosomal segments with a power-law function fitted to the data (blue line). Data are colored by their entanglement value (rainbow color palette). (**D**) Distribution and variation of the entanglement score (ES) of the two chromosomal segments being compared. (**B and D**) Box boundaries represent 1st and 3rd quartiles, middle line represents median, and whiskers extend to 1.5 times the interquartile range, (Mann-Whitney rank, *: p < 5 x 10^−2^). (**E, F**) Hierarchical clustering of (**E**) mean distance scores (MDS) and (**F**) mean entanglement scores (MES) reveal two major groups of chromosomal segments.

**Fig. S6.**
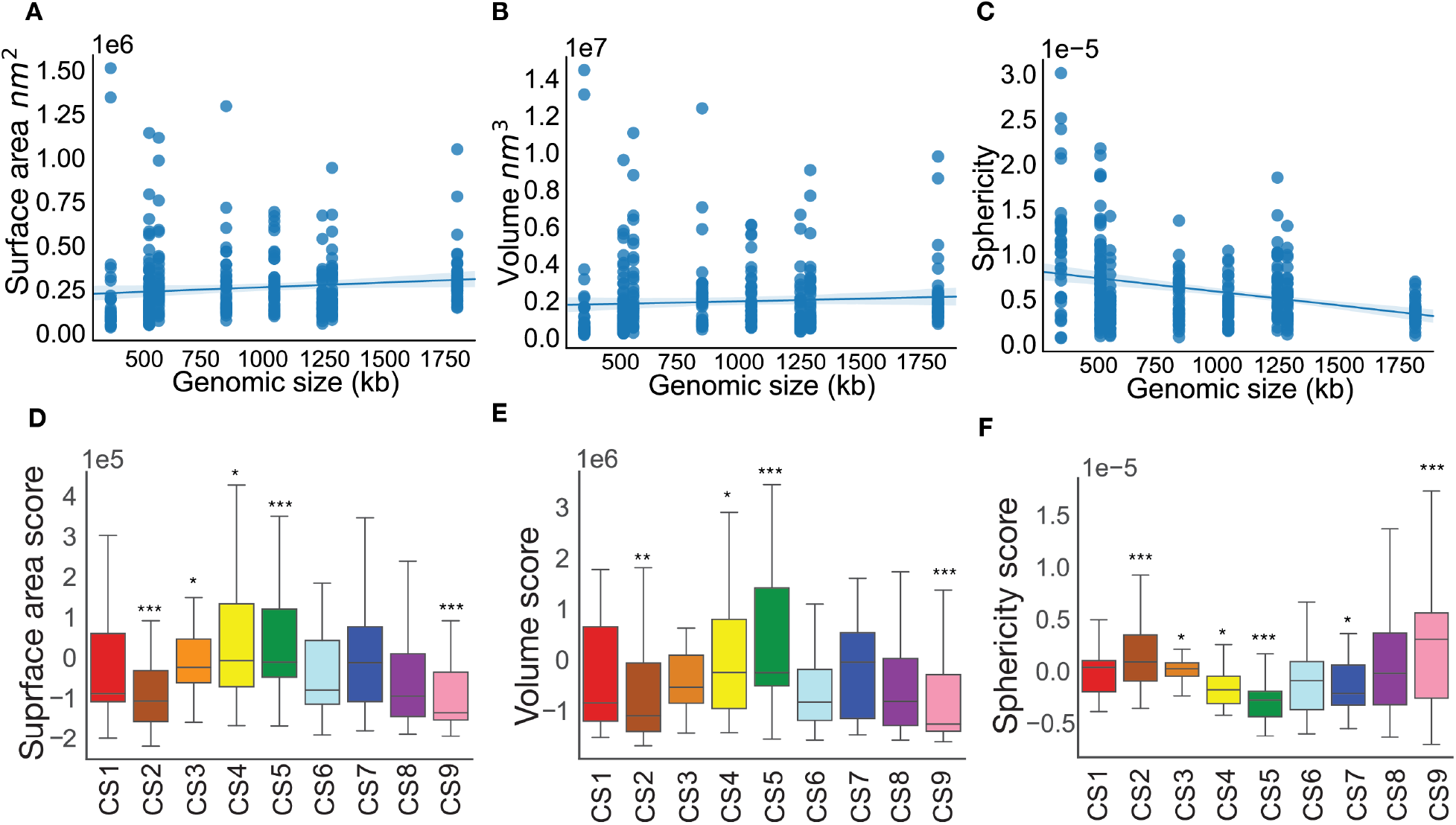
3D features of chromosomal segments. (**A-C**) Surface area, volume and sphericity versus genomic distance for the 342 analyzed chromosomal segments (linear fit in blue). (**D-F**) Box plots showing distribution and variation of the 9 chromosomal segments in terms of (**D**) surface area, (**E**) volume, and (**F**) sphericity score. The chromosomal segments are color-coded as in Figure 2A. Box boundaries represent 1st and 3rd quartiles, middle line represents median, and whiskers extend to 1.5 times the interquartile range (Mann-Whitney rank test, *: p < 5 x 10^−2^, **: p < 10^−2^, ***: p < 10^−3^).

**Fig. S7.**
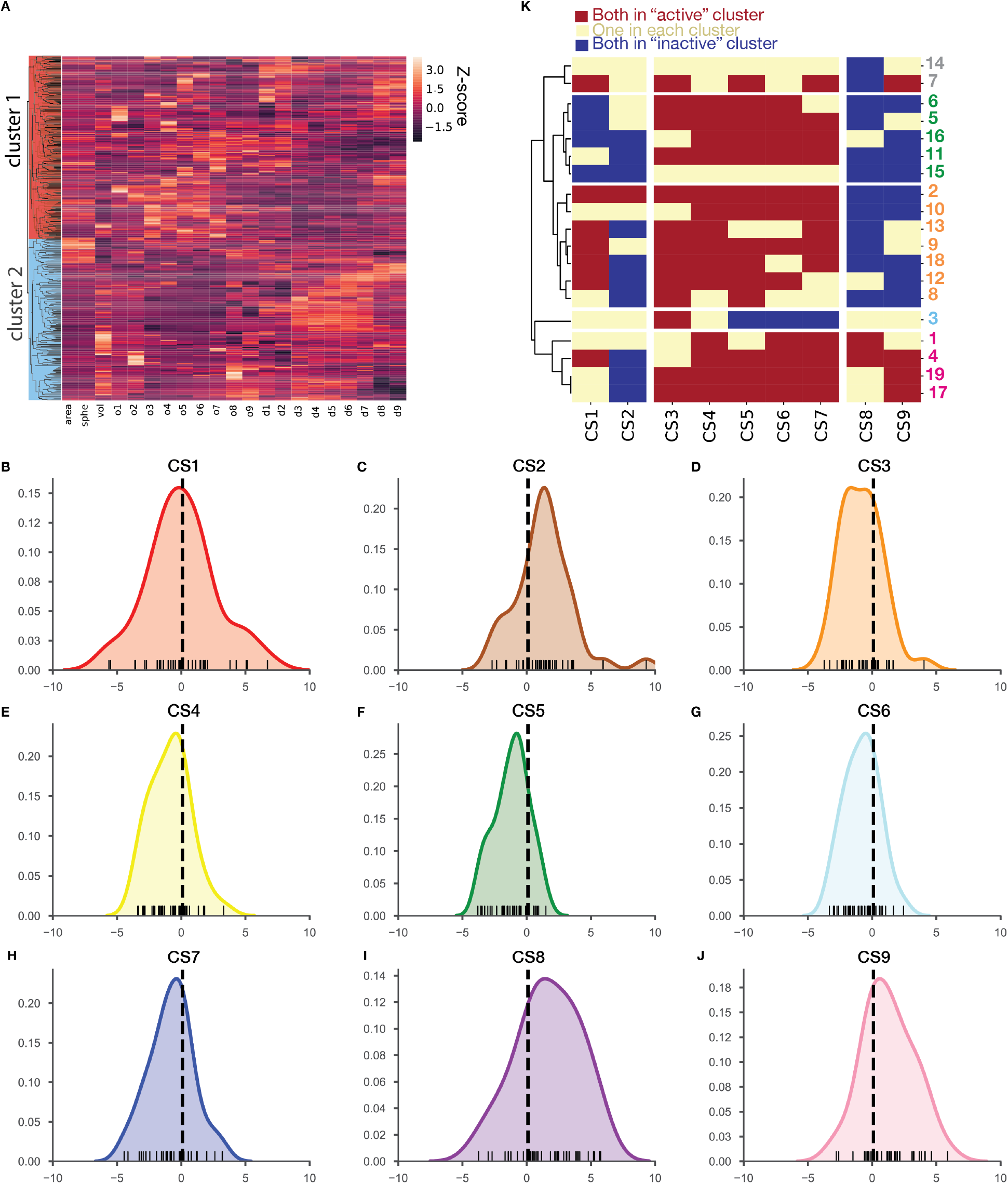
Clustering of chromosomal segments. (**A**) Unsupervised hierarchical clustering of 342 analyzed chromosomal segments, arranged on the y-axis by Euclidian distance (dendrogram). Colors on the heat map represent the Z-score of the 3D features [surface area, volume, and sphericity scores (o1-o9), MDS, and MES (d1-d9)] for each chromosomal segment. Clustering reveals two major groupings of chromosomal segments. (**B-J**) Distribution along PC1 for CS1-9. Color-coded as in Figure 2. Vertical black dotted line represents the partition in cluster 1 and cluster 2. (**K**) Hierarchical clustering of the compartment state profile vectors for the 19 analyzed nuclei results in 5 large clusters. The compartment state profile (Methods) was calculated comparing the chromatin state profile of each homolog within a single nucleus. Note, the two most populated clusters confirm that, in the majority of the analysed nuclei, the compartment state profile is maintained with CS2, 8, and 9 mostly inactive, CS3-7 mostly active, and CS1seeming to occupy both states.

**Fig. S8.**
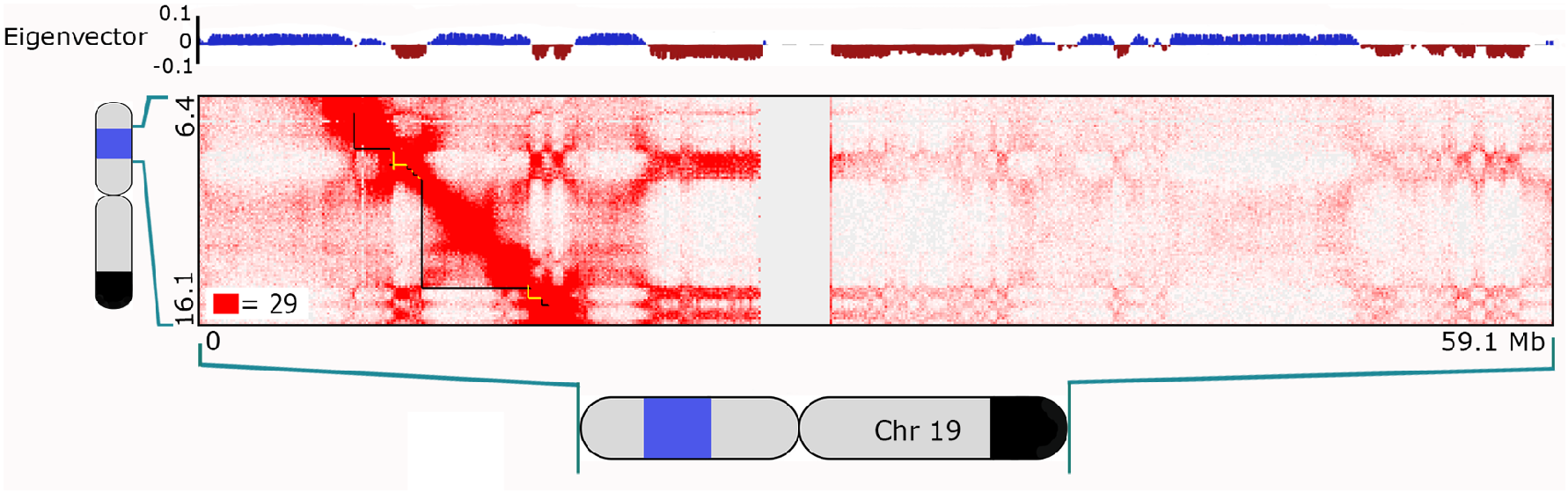
*In situ* Hi-C analysis of 8.16 Mb imaged region segregates into two compartments. Map shows region including CS1-9, displayed at 100kb resolution and normalized using Knight & Ruiz (51). Contact count indicated by the color of each pixel, ranging from 0 (white) to 29 (red); manually curated intervals in compartment A and B are delineated below the diagonal in black and yellow, respectively, and reflect the positive values (blue) and negative values (brown), respectively, of the compartment eigenvector at 100 kb resolution (above map). Cartoons of chromosome 19 show extent of the 8.16 Mb imaged region (blue) and block of SNVs shared between PGP1 and PGP95 (black; as in Fig. 1B).

**Fig. S9.**
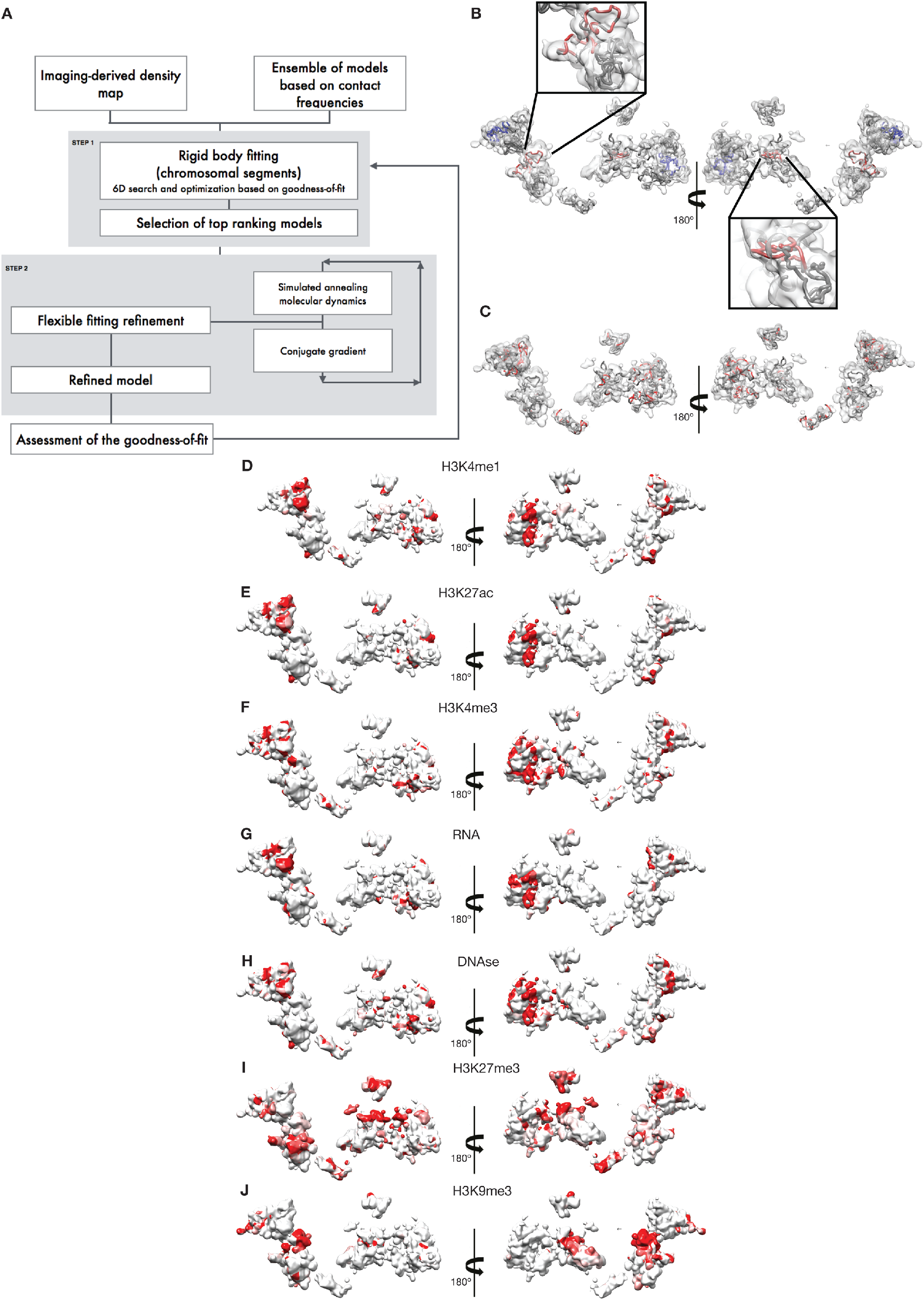
Genomic features of the modeled 8.16 Mb region. (**A**) IMGR protocol for fitting and refinement of 3D chromatin models. The protocol inputs are a density map derived from a single PGP1f nucleus imaged with OligoSTORM and an ensemble of 3D models derived from a PGP1f population-based *in situ* Hi-C contact map. The protocol entails rigid fitting by a random 6D search (step1), followed by flexible fitting refinement via simulated annealing rigid-body molecular dynamics and conjugate gradient minimization (step 2). After each step, the goodness-of-fit is assessed as described using different scores (Supplementary note). (**B**) Mapping of Znf clusters onto the 3D trace. The trace is colored in grey, and the two clusters are colored in blue and red. Insets are zoomed-in views of one of the Znf clusters, highlighting the differences in conformation of the cluster in the model of the two homologs. (**C**) Annotated disease-causing genes mapped into the 3D trace. The trace is colored in grey. Shown in red are genes for which the molecular basis for the disorder is known and a mutation has been found, as classified by OMIM Genes track in the UCSC Genome Browser (92). (**D-J**) Mapping of genome features. Genomic features are presented as mean value per 10kb. The color scale for each genomic feature ranges from white (mean value) to red (mean + 2 standard deviation). The figures in this panel were created with Chimera (50).

**Fig. S10.**
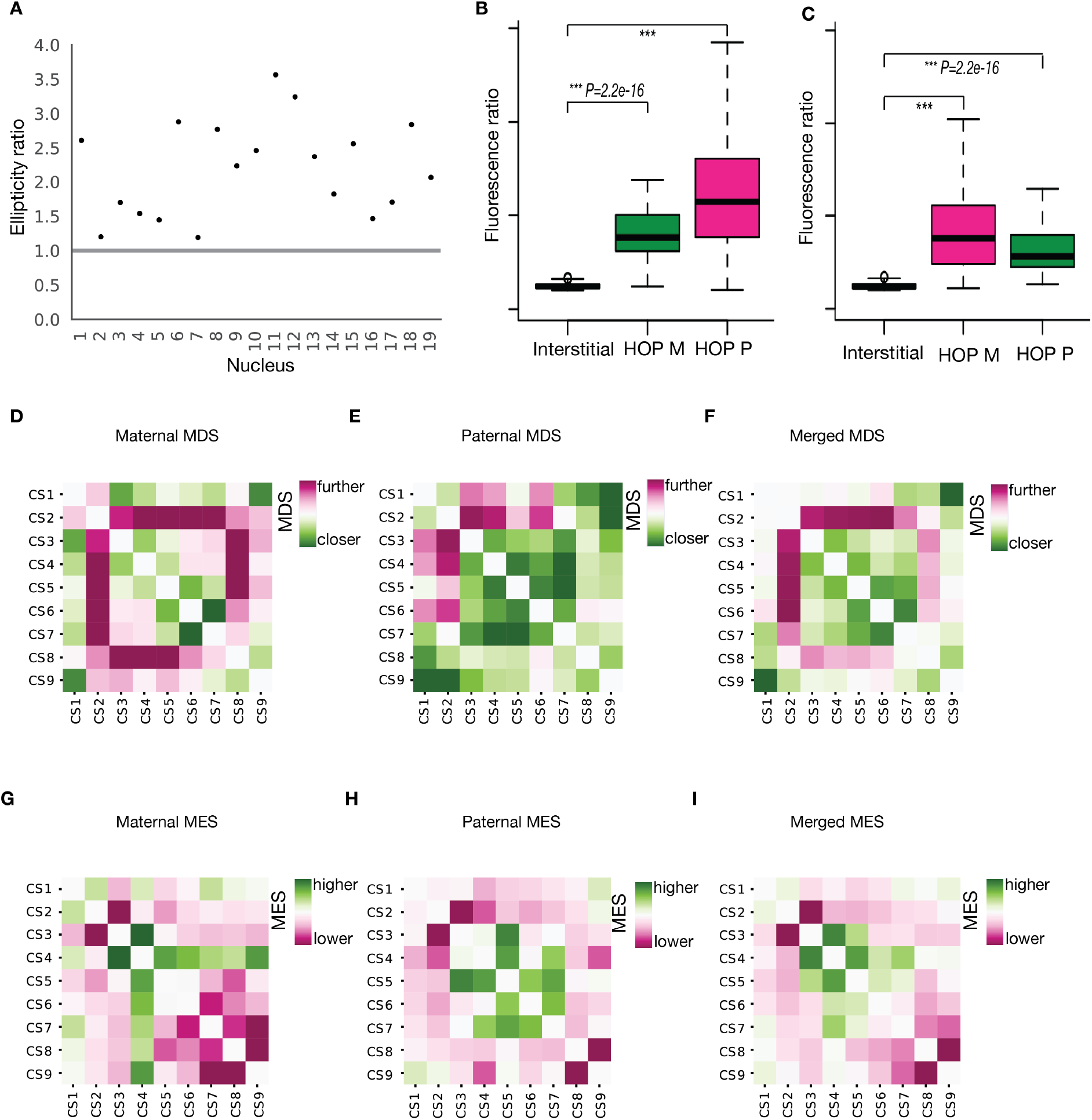
Ellipticity ratios and homolog-specific analyses. (**A**) Ellipticity ratios of the 19 nuclei analyzed (see Table S6 for values of ellipticity ratios). Grey line, ellipticity score (1.0) expected when the two largest principal axes are equal. (**B**) Ratios of signal intensity using interstitial, HOP-M (Atto488N), and HOP-P (Atto565N) probes. (**C**) Same as B, except HOP-M is now labeled with Atto565N, and HOP-P is labeled with Atto488N. (**D-I**) The MDS and MES matrices for six nuclei showing only (**D, G**) maternal homologs, (**E, H**) only paternal homologs, and (**F, I**) maternal and paternal homologs, combined.

**Fig. S11.**
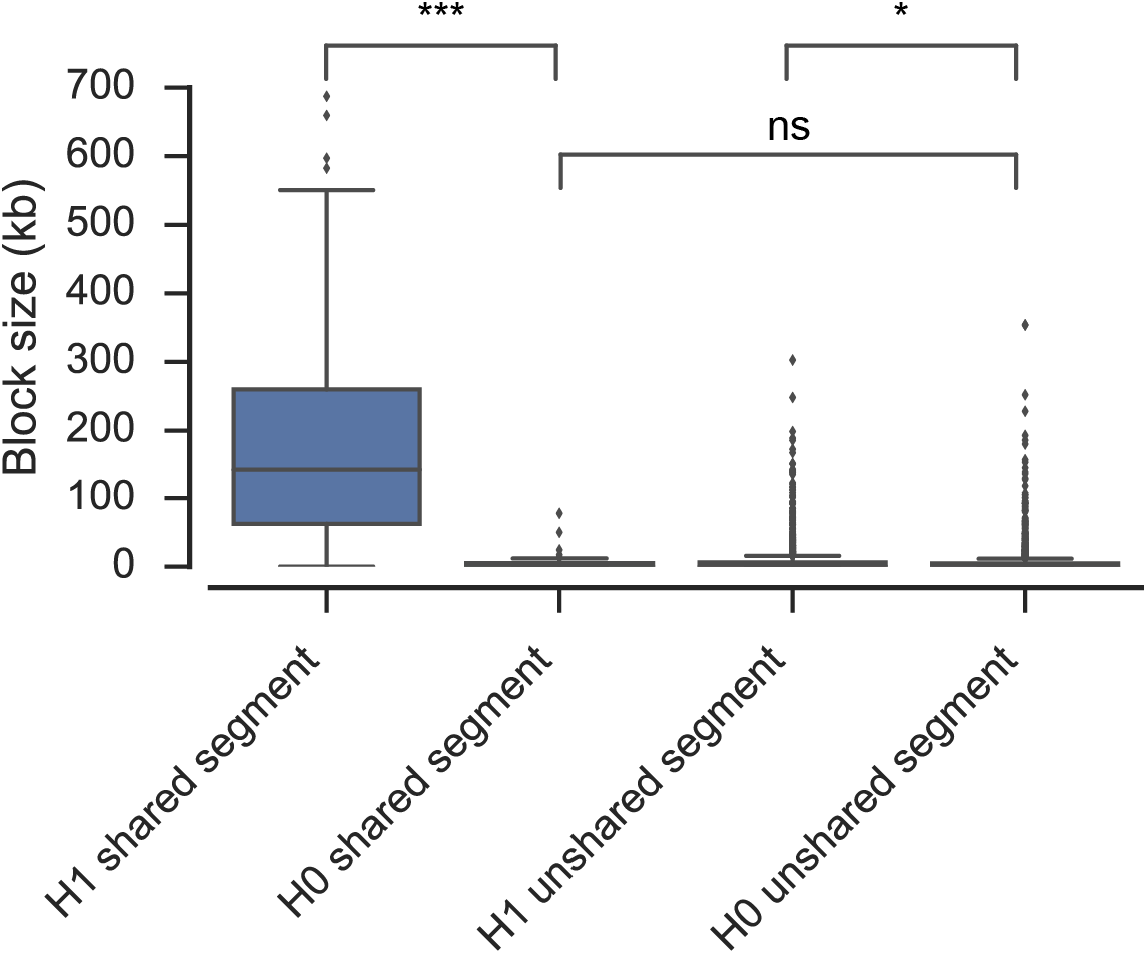
Identification of the maternally derived haplotype of PGP1. The longest blocks of SNVs on chromosome 19 of PGP1 that are shared with PGP95 correspond to the H1 haplotype, thus identifying homologs bearing long blocks of H1 SNVs as maternally derived (Mann-Whitney rank test, ns: non-significant, *: p < 5 x 10^−2^, ***: p < 10^−3^).

**Fig. S12.**
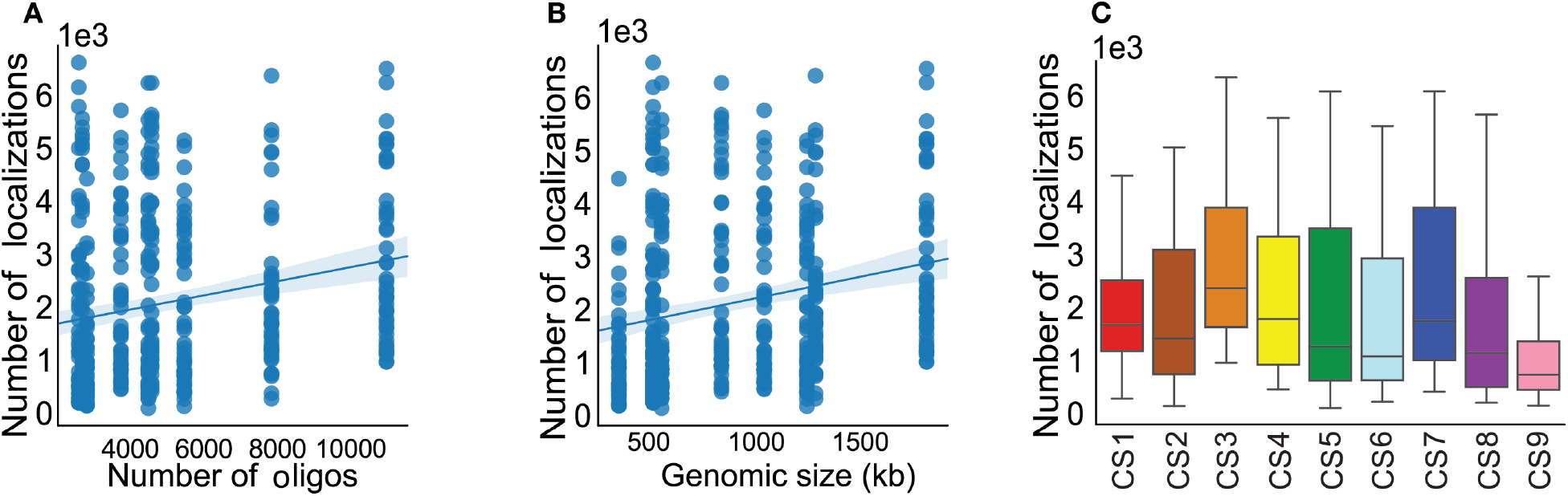
(**A**) Number of localizations versus number of oligos and (**B**) genomic distance for the 342 analyzed chromosomal segments (linear fit in blue). (**C**) Number of localizations per round of imaging. Color coding as in Figure S7.

**Fig. S13.**
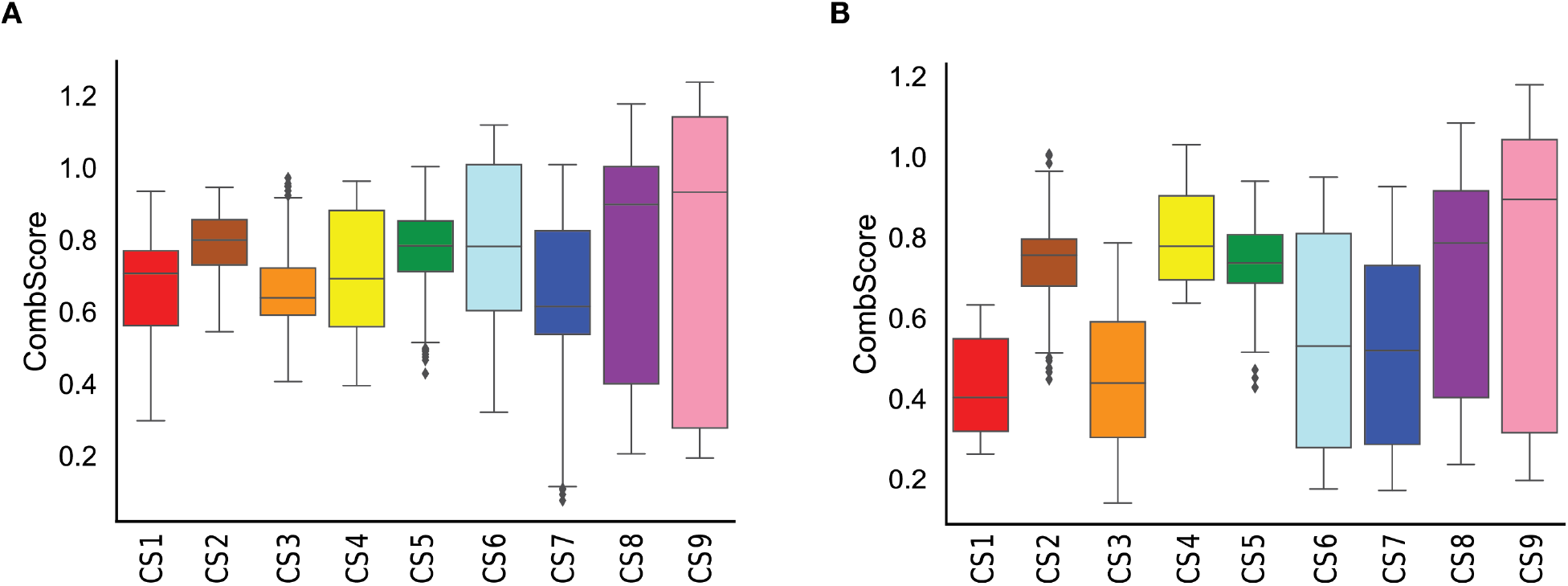
Assessment of the rigid body step of the fitting protocol. (**A, B**) Distribution of the Combined Score (CombScore) for each chromosomal segment of each of the two homologs traced. The chromosomal segments are color-coded as in Figure 2A and B.

**Fig. S14.**
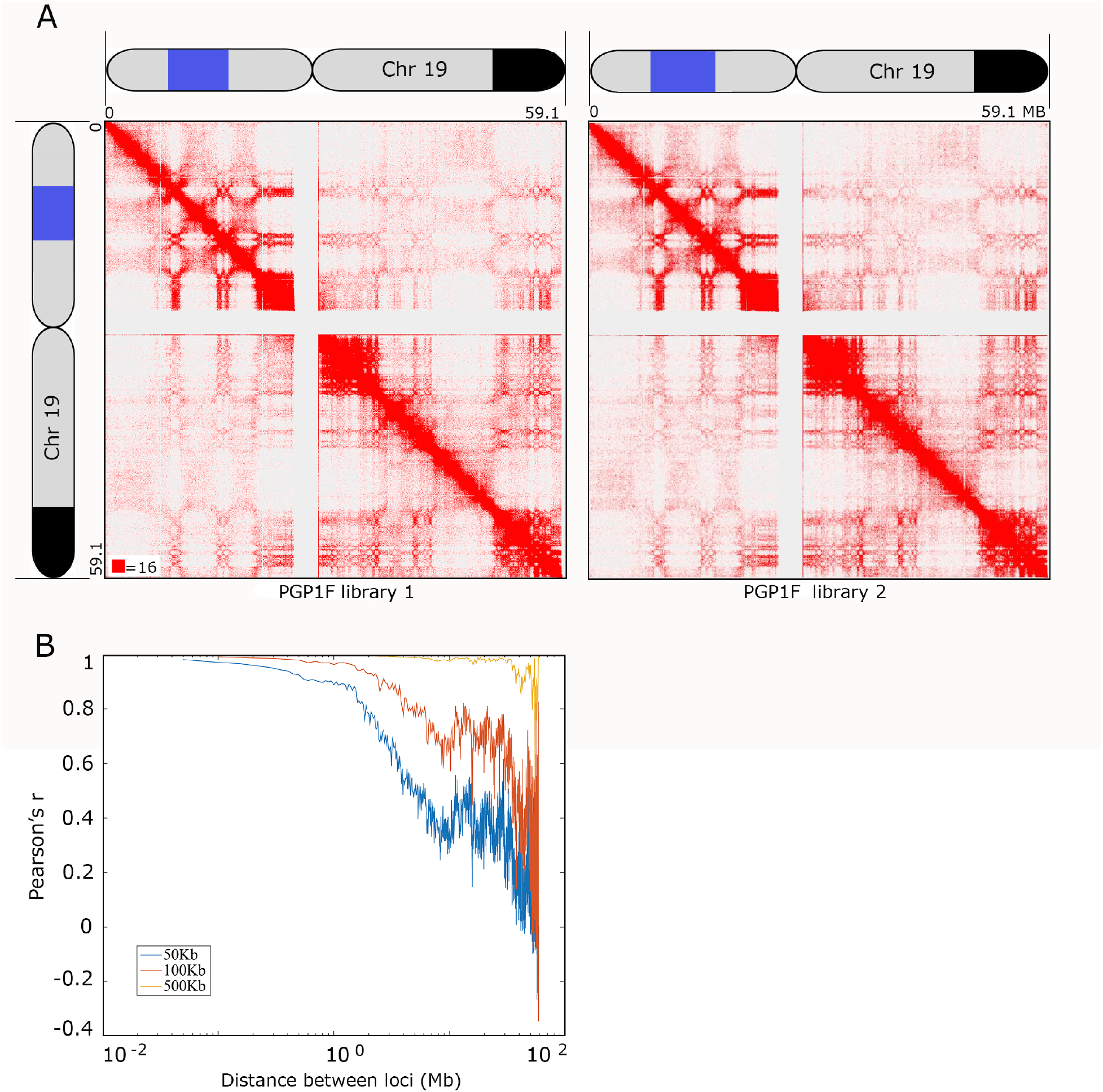
*In situ* Hi-C results for PGP1f are reproducible. (**A**) The outcomes of two Hi-C experiments conducted on PGP1f cells are extremely similar, as shown here and in Table S9 The Hi-C maps represent the entirety of chromosome 19, displayed at 100 kb resolution and normalized according to Knight & Ruiz (51). Contact count indicated by the color of each pixel, ranging from 0 (white) to 16 (red). Cartoons of chromosome 19 show extent of the 8.16 Mb imaged region (blue) and block of SNVs shared between PGP1 and PGP95 (black; as in Fig. 1B). (**B**) Correlation as a function of distance between *in-situ* Hi-C PGP1f libraries 1 and 2; 50 (blue line), 100 (mahogany line) and 500 kb (yellow line) resolutions are shown. Hi-C replicates are highly correlated as demonstrated by the Pearson’s r value; for distances of tens of megabases, r> 0.95 at 500 kb; for 1 Mb distance, r> 0.96 at 100 kb and r=0.89 at 50 kb. Our results indicate high similarity between PGP1f *in-situ* Hi-C duplicates.

### Tables

**Table S1.** Chromosomal segments imaged with OligoSTORM.

| Round | Name | Start<br>(hg19) | End<br>(hg19) | Size<br>(kb) | Number<br>of<br>oligos | Average # of<br>localizations/<br>Cluster* |
| --- | --- | --- | --- | --- | --- | --- |
| 1 | CS1 | 7,400,000 | 86,80,000 | 1,280 | 7,861 | 4,825 |
| 2 | CS2 | 8,680,000 | 9,920,000 | 1,240 | 5,489 | 2,979 |
| 3 | CS3 | 9,920,000 | 11,720,000 | 1,800 | 10,985 | 4,489 |
| 4 | CS4 | 11,720,000 | 12,760,000 | 1,040 | 3,768 | 2,759 |
| 5 | CS5 | 12,760,000 | 13,320,000 | 560 | 4,516 | 2,560 |
| 6 | CS6 | 13,320,000 | 13,840,000 | 520 | 2,720 | 1,861 |
| 7 | CS7 | 13,840,000 | 14,680,000 | 840 | 4,601 | 2,620 |
| 8 | CS8 | 14,680,000 | 15,200,000 | 520 | 2,624 | 2,068 |
| 9 | CS9 | 15,200,000 | 15,560,000 | 360 | 2,843 | 1,389 |
| 10 | CD51 | 13,840,000 | 13,980,000 | 140 | 751 | 1,045 |
| 11 | CD52 | 13,980,000 | 14,240,000 | 260 | 1,698 | 1,526 |
| 12 | CD53 | 14,240,000 | 14,590,000 | 350 | 1,639 | 1,760 |
| 13 | CD54 | 14,590,000 | 14,680,000 | 90 | 513 | 516 |
| 14 | Loop anchor<br>in CS 6 | 13,400,000 | 13,410,000 | 10 | 41 | 32 |
| 15 | Loop body in CS 6 | 13,410,001 | 13,680,000 | 270 | 1,473 | 134 |
| 16 | Loop anchor<br>in CS 6 | 13,680,001 | 13,690,000 | 10 | 28 | 99 |
| 17 | Downstream<br>loop flank in<br>CS 6 | 13,690,000 | 13,710,000 | 20 | 104 | 46 |
| 18 | Upstream<br>loop flank in<br>CS 6 | 13,320,000 | 13,400,000 | 80 | 604 | 134 |
| 19 | *DNMT1*<br>gene | 10,244,022 | 10,303,505 | 59.483 | 529 | 42 |

**Table S2.** Multiple chromosome walk.

A.
| **Round** | **Start (hg19)** | **End (hg19)** | **Size**<br>**(kb)** | **Number**<br>**of**<br>**oligos** | **Average # of**<br>**localizations/cluster\*** |
| --- | --- | --- | --- | --- | --- |
| 5 | 120,000,000 | 120,250,000 | 250 | 864 | 558 |
| 6 | 120,250,000 | 120,500,000 | 250 | 989 | 652 |
| 7 | 120,500,000 | 120,750,000 | 250 | 684 | 513 |
| 8 | 120,750,000 | 121,000,000 | 250 | 547 | 198 |

B.
| **Round** | **Start (hg19)** | **End (hg19)** | **Size**<br>**(kb)** | **Number**<br>**of**<br>**oligos** | **Average # of**<br>**localizations/cluster\*** |
| --- | --- | --- | --- | --- | --- |
| 6 | 150,000,000 | 150,500,000 | 500 | 2,524 | 1,503 |
| 7 | 150,500,000 | 151,000,000 | 500 | 3,467 | 2,320 |
| 8 | 151,000,000 | 151,500,000 | 500 | 2,511 | 878 |

**(A)** Walk along chromosome 5. **(B)** Walk along chromosome 3.
\* As in Figure E.

**Table S3.** Chromosomal segments imaged with OligoDNA-PAINT.

| **Segment** | **Imager** | **Imager conc. (nM)** | **# of localizations\*** | **Median fit precision (nm)** | **NeNA localization precision in XY (nm)** | **Supported resolution in XY (nm)** | **On (s)** | **Off (s)** |
| --- | --- | --- | --- | --- | --- | --- | --- | --- |
| CS7 | P1 | 1 | 5,462 | 4.30 | 9.43 | 22.15 | 3.11 | 10.85 |
| CS8 | P13 | 1 | 6,973 | 2.96 | 8.00 | 18.79 | 2.64 | 8.75 |
| CS9 | P9 | 1 | 11,348 | 2.33 | 5.40 | 12.68 | 3.45 | 7.85 |

\* As shown in Figure 2F
Number of frames for each segment was 30,000.

**Table S4.** Compartment classification

| CS | Active | Inactive | Hi-C classification |
| --- | --- | --- | --- |
| 4 | 87 | 13 | A |
| 3 | 84 | 16 | A |
| 5 | 84 | 16 | A |
| 7 | 82 | 18 | A |
| 6 | 79 | 21 | A |
| 1 | 58 | 42 | A |
| 9 | 37 | 63 | A |
| 2 | 26 | 72 | B |
| 8 | 24 | 76 | B |

Chromosomal segment (CS); Active, % active CS; Inactive, % inactive CS; Hi-C classification (see Method).

**Table S5.** Hi-C library statistics for PGP1F “mega map” (combined data from library 1 and library 2).

|  | **Combined data from Library 1 and Library 2** |
| --- | --- |
| **Sequenced Read Pairs (Status)** | 691,620,560 |
| **Normal Paired** | 360,608,028 (52.14%) |
| **Chimeric Paired** | 273,227,134 (39.51%) |
| **Chimeric Ambiguous** | 45,051,055 (6.51%) |
| **Unalignable** | 12,734,343 (1.84%) |
| **Ligation Motif Present** | 446,227,577 (64.52%) |
| **Alignable (Normal+Chimeric Paired)** | 633,835,162 (91.64%) |
| **Unique Read Pairs** | 597,056,142 (86.33%) |
| **PCR Duplicates** | 34,709,496 (5.02%) |
| **Optical Duplicates** | 2,069,524 (0.30%) |
| **Library Complexity Estimate** | 5,536,972,413 |
| **Intra-fragment Read Pairs** | 5,697,684 (0.82% / 0.95%) |
| **Below MAPQ Threshold** | 52,118,887 (7.54% / 8.73%) |
| **Hi-C Contacts** | 539,239,571 (77.97% / 90.32%) |
| **Ligation Motif Present** | 238,907,673 (34.54% / 40.01%) |
| **3' Bias (Long Range)** | 72% - 28% |
| **Pair Type % (L-I-O-R)** | 25% - 25% - 25% - 25% |
| **Inter-chromosomal** | 129,868,605 (18.78% / 21.75%) |
| **Intra-chromosomal** | 409,370,966 (59.19% / 68.56%) |
| **Short Range (<20Kb)** | 112,473,822 (16.26% / 18.84%) |
| **Long Range (>20Kb)** | 296,896,731 (42.93% / 49.73%) |

**Table S6.**
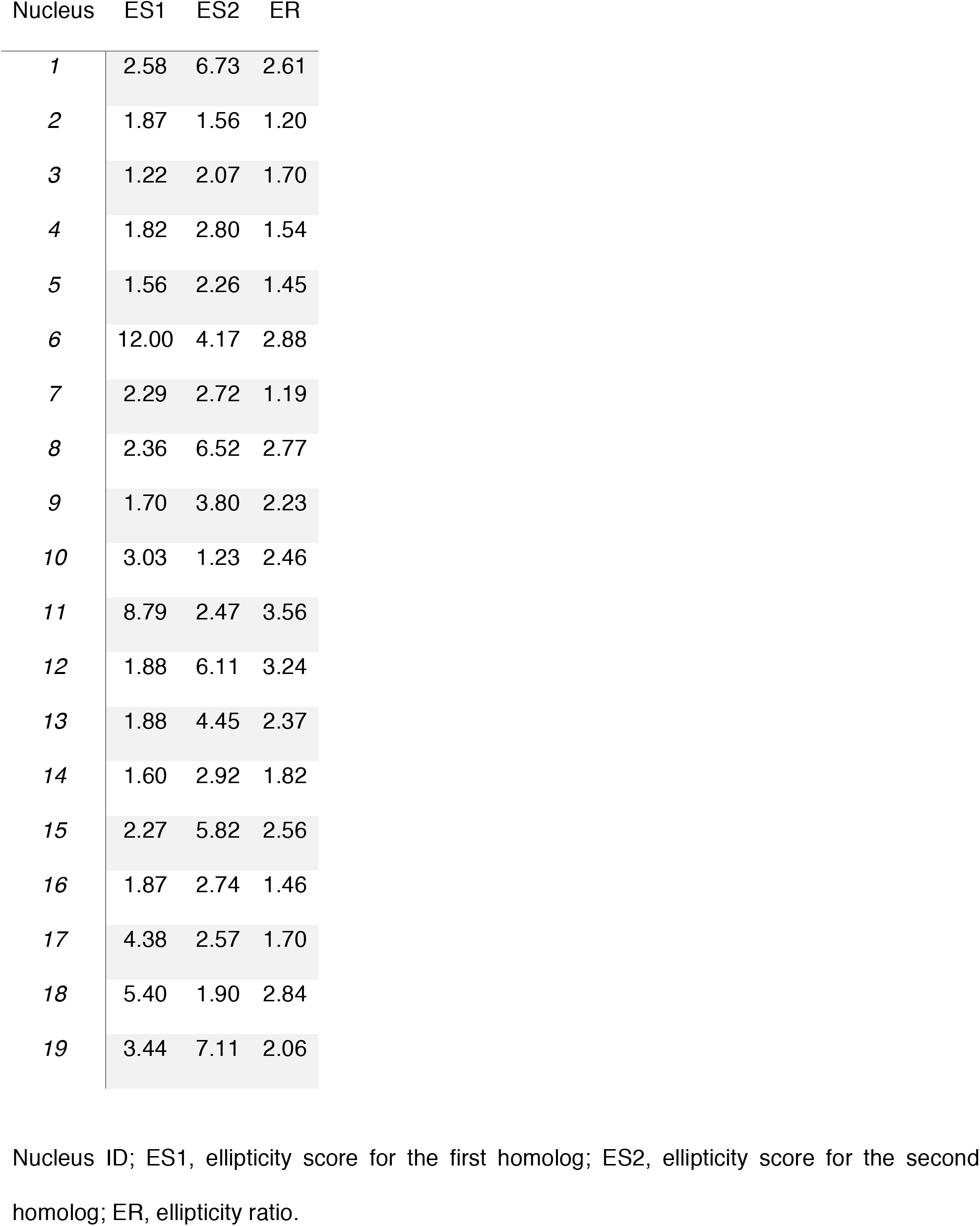
Homologs Ellipticity in the cell population.

| Nucleus | ES1 | ES2 | ER |
| --- | --- | --- | --- |
| 1 | 2.58 | 6.73 | 2.61 |
| 2 | 1.87 | 1.56 | 1.20 |
| 3 | 1.22 | 2.07 | 1.70 |
| 4 | 1.82 | 2.80 | 1.54 |
| 5 | 1.56 | 2.26 | 1.45 |
| 6 | 12.00 | 4.17 | 2.88 |
| 7 | 2.29 | 2.72 | 1.19 |
| 8 | 2.36 | 6.52 | 2.77 |
| 9 | 1.70 | 3.80 | 2.23 |
| 10 | 3.03 | 1.23 | 2.46 |
| 11 | 8.79 | 2.47 | 3.56 |
| 12 | 1.88 | 6.11 | 3.24 |
| 13 | 1.88 | 4.45 | 2.37 |
| 14 | 1.60 | 2.92 | 1.82 |
| 15 | 2.27 | 5.82 | 2.56 |
| 16 | 1.87 | 2.74 | 1.46 |
| 17 | 4.38 | 2.57 | 1.70 |
| 18 | 5.40 | 1.90 | 2.84 |
| 19 | 3.44 | 7.11 | 2.06 |

Nucleus ID; ES1, ellipticity score for the first homolog; ES2, ellipticity score for the second homolog; ER, ellipticity ratio.

**Table S7.**
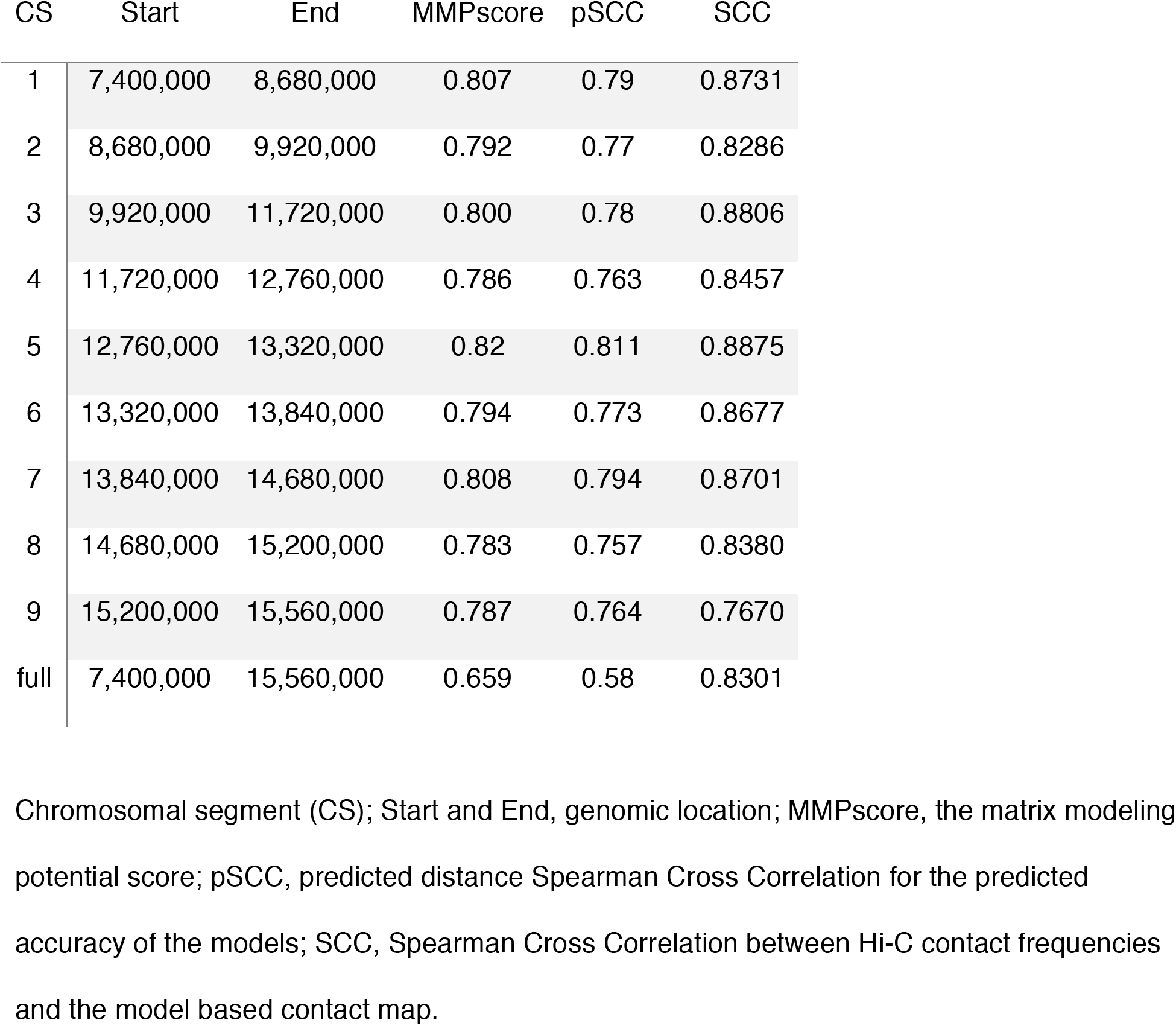
Assessment of potential for modeling of the analyzed regions.

| CS | Start | End | MMPscore | pSCC | SCC |
| --- | --- | --- | --- | --- | --- |
| 1 | 7,400,000 | 8,680,000 | 0.807 | 0.79 | 0.8731 |
| 2 | 8,680,000 | 9,920,000 | 0.792 | 0.77 | 0.8286 |
| 3 | 9,920,000 | 11,720,000 | 0.800 | 0.78 | 0.8806 |
| 4 | 11,720,000 | 12,760,000 | 0.786 | 0.763 | 0.8457 |
| 5 | 12,760,000 | 13,320,000 | 0.82 | 0.811 | 0.8875 |
| 6 | 13,320,000 | 13,840,000 | 0.794 | 0.773 | 0.8677 |
| 7 | 13,840,000 | 14,680,000 | 0.808 | 0.794 | 0.8701 |
| 8 | 14,680,000 | 15,200,000 | 0.783 | 0.757 | 0.8380 |
| 9 | 15,200,000 | 15,560,000 | 0.787 | 0.764 | 0.7670 |
| full | 7,400,000 | 15,560,000 | 0.659 | 0.58 | 0.8301 |

Chromosomal segment (CS); Start and End, genomic location; MMPscore, the matrix modeling potential score; pSCC, predicted distance Spearman Cross Correlation for the predicted accuracy of the models; SCC, Spearman Cross Correlation between Hi-C contact frequencies and the model based contact map.

**Table S8.** Flexible fitting refinement: assessment of the goodness-of-fit.

| CS | CCCs |  | CCCf |  | CLSs |  | CLSf |  | RMSDs-f |  |
| --- | --- | --- | --- | --- | --- | --- | --- | --- | --- | --- |
| 1 | 0.47 | 0.59 | 0.61 | 0.83 | 0.17 | 0.40 | 0.20 | 0.50 | 14.32 | 19.13 |
| 2 | 0.57 | 0.53 | 0.72 | 0.76 | 0.45 | 0.26 | 0.55 | 0.30 | 16.20 | 22.23 |
| 3 | 0.50 | 0.42 | 0.74 | 0.70 | 0.60 | 0.58 | 0.66 | 0.65 | 20.90 | 22.76 |
| 4 | 0.64 | 0.60 | 0.77 | 0.71 | 0.34 | 0.36 | 0.40 | 0.45 | 19.43 | 10.35 |
| 5 | 0.62 | 0.53 | 0.69 | 0.78 | 0.14 | 0.10 | 0.16 | 0.13 | 13.90 | 19.27 |
| 6 | 0.68 | 0.60 | 0.78 | 0.81 | 0.16 | 0.16 | 0.21 | 0.17 | 16.30 | 16.30 |
| 7 | 0.66 | 0.55 | 0.82 | 0.84 | 0.25 | 0.25 | 0.32 | 0.35 | 18.89 | 17.29 |
| 8 | 0.70 | 0.63 | 0.82 | 0.82 | 0.20 | 0.19 | 0.22 | 0.24 | 15.20 | 17.42 |
| 9 | 0.78 | 0.72 | 0.85 | 0.90 | 0.12 | 0.12 | 0.17 | 0.18 | 16.00 | 14.48 |

Chromosomal segment (CS); Cross Correlation Coefficient of the starting rigid fitted model (CCCs); Cross Correlation Coefficient of the final model after flexible fitting refinement (CCCf); Clash Score of the starting rigid fitted model (CLSs); Clash Score of the final model after flexible fitting refinement (CLSf); Root Mean Square Deviation between starting rigid fitted model and final model after flexible fitting refinement (RMSDs-f).

**Table 9.** Hi-C library statistics for PGP1F replicates.

|  | **Library 1** | **Library 2** |
| --- | --- | --- |
| **Sequenced Read Pairs**<br>**(Status)** | 299,490,370 | 392,130,190 |
| **Normal Paired** | 169,241,991 (56.51%) | 191,366,037<br>(48.80%) |
| **Chimeric Paired** | 102,483,082 (34.22%) | 170,744,052<br>(43.54%) |
| **Chimeric Ambiguous** | 19,764,751 (6.60%) | 25,286,304<br>(6.45%) |
| **Unalignable** | 8,000,546 (2.67%) | 4,733,797<br>(1.21%) |
| **Ligation Motif Present** | 172,812,075 (57.70%) | 273,415,502<br>(69.73%) |
| **Alignable**<br>**(Normal+Chimeric**<br>**Paired)** | 271,725,073 (90.73%) | 362,110,089<br>(92.34%) |
| **Unique Read Pairs** | 258,780,678 (86.41%) | 339,925,404<br>(86.69%) |
| **PCR Duplicates** | 12,074,226 (4.03%) | 20,980,766<br>(5.35%) |
| **Optical Duplicates** | 870,169 (0.29%) | 1,203,919<br>(0.31%) |
| **Library Complexity Estimate** | 2,947,002,205 | 2,982,605,992 |
| **Intra-fragment Read Pairs** | 2,212,105 (0.74% / 0.85%) | 3,528,694<br>(0.90% / 1.04%) |
| **Below MAPQ Threshold** | 22,946,825 (7.66% / 8.87%) | 29,492,420<br>(7.52% / 8.68%) |
| **Hi-C Contacts** | 233,621,748 (78.01% / 90.28%) | 306,904,290<br>(78.27% / 90.29%) |
| **Ligation Motif Present** | 92,975,257 (31.04% / 35.93%) | 147,078,375<br>(37.51% / 43.27%) |
| **3' Bias (Long Range)** | 69% - 31% | 74% - 26% |
| **Pair Type % (L-I-O-R)** | 25% - 25% - 25% - 25% | 25% - 25% - 25%<br>- 25% |
| **Inter-chromosomal** | 55,054,421 (18.38% / 21.27%) | 74,848,460<br>(19.09% / 22.02%) |
| **Intra-chromosomal** | 178,567,327 (59.62% / 69.00%) | 232,055,830<br>(59.18% / 68.27%) |
| **Short Range (<20Kb)** | 48,073,771 (16.05% / 18.58%) | 65,627,376<br>(16.74% / 19.31%) |
| **Long Range (>20Kb)** | 130,493,315 (43.57% / 50.43%) | 166,428,281<br>(42.44% /<br>48.96%) |

**Movie S1**. OligoSTORM imaging of the 8.16 Mb region of human chromosome 19 encompassing CS1-9. Nucleus is the same as that shown in Figure 2B. Color coding as in Figure 2A, B.

**Movie S2**. Model obtained via IMGR of the 8.16 Mb region of human chromosome 19 encompassing CS1-9. Nucleus is the same as that shown in Figure 2B. Color coding as in Figure 2A, B.

